# Computing Absolute Binding Affinities by Streamlined Alchemical Free Energy Perturbation (SAFEP)

**DOI:** 10.1101/2022.12.09.519809

**Authors:** Ezry Santiago-McRae, Mina Ebrahimi, Jesse W. Sandberg, Grace Brannigan, Jérôme Hénin

## Abstract

Free Energy Perturbation (FEP) is a powerful but challenging computational technique for estimating differences in free energy between two or more states. This document is intended both as a tutorial and as an adaptable protocol for computing free energies of binding using free energy perturbations in NAMD. We present the Streamlined Alchemical Free Energy Perturbation (SAFEP) framework. SAFEP shifts the computational frame of reference from the ligand to the binding site itself. This both simplifies the thermodynamic cycle and makes the approach more broadly applicable to superficial sites and other less common geometries. As a practical example, we give instructions for calculating the absolute binding free energy of phenol to lysozyme. We assume familiarity with standard procedures for setting up, running, and analyzing molecular dynamics simulations using NAMD and VMD. While simulation times will vary, the human tasks should take no more than 3 to 4 hours for a reader without previous training in free energy calculations or experience with the VMD Colvars Dashboard. Sample data are provided for all key calculations both for comparison and readers’ convenience.

## 1 Introduction

In this tutorial, we are principally concerned with computing the Absolute Binding Free Energy (ABFE) of a ligand to its receptor. While many methods of measuring free energies exist, alchemical Free Energy Perturbation (FEP) methods make use of the fact that since the change in free energy is path-independent it can be calculated via an unphysical path. In the case of FEP, that unphysical path is defined by scaling the non-bonded interactions of the ligand [1–6]. In essence, the user can make a bound ligand “disappear” from the binding site, make it re-appear in the bulk solution, and calculate the corresponding free energy difference.

While elegant and exact in principle, FEP calculations are often unwieldy in practice. One of the most stubborn challenges that most FEP implementations face is that the ligand must maintain the original bound configuration during decoupling, even as the very interactions that stabilize the bound configuration are weakened. Consequently, absolute binding schemes introduce restraints on the ligand to mimic the interactions that stabilize the bound ensemble. While there are many such schemes to accelerate convergence, [7– 10] most do so in ways that require error-prone manual parameterization and many hours of the user’s time. Many of these techniques have already been reviewed by Mey et al [2]. As discussed there, no restraint scheme is without cost; the restraints themselves will bias the estimated free energy and must be corrected in the coupled state, the decoupled (gas phase) state, or both. Conveniently, restraints that have simple geometries *and* do not affect the coupled state can be accounted for analytically. On the other hand, numerical approaches must be used to correct for restraints (such as an RMSD [10] or DBC collective variable [11]) that restrict the accessible configuration space to an irregular volume. However, if the restraint potential on the collective variable is chosen to have negligible impact on the coupled state (such as a flat-well potential [12]), these calculations can be done cheaply in vacuum as illustrated in this tutorial. When more intrusive restraints (such as harmonic potentials [9, 10]) are applied in the coupled state, it is necessary to calculate numerical corrections in the coupled state regardless of the restraint geometry.

Streamlined Alchemical Free Energy Perturbation (SAFEP) is specifically designed to make FEP calculations faster and easier for the user without sacrificing the accuracy of the final free energy estimate. SAFEP reduces conceptual and computational complexity by replacing conventional rotational and translational restraints for stabilizing the ligand in the binding site with a single Distance-to-Bound Configuration (DBC) coordinate. This makes setup easier because instead of parameterizing six or more individual restraints, the user need only parameterize one. Furthermore SAFEP can handle superficial binding sites in phase-separated bulk with atomistic resolution [11]. Statistically optimal FEP estimators require both decoupling and recoupling calculations; SAFEP uses Interleaved Double-Wide Sampling (IDWS) to extract both quantities from the same calculation, roughly halving the required simulation time (see the `alchLambdaIDWS` entry in the [NAMD User Guide section on Free Energy Perturbation](https://www.ks.uiuc.edu/Research/namd/2.14/ug/node64.html)) [13]. SAFEP makes extensive use of the Colvars Dashboard in VMD allowing the user to easily measure collective variables, impose biases, and generate restraint configuration files from one interface [14]. Finally, analysis tools and data visualizations are included in one Jupyter notebook allowing for comprehensive quality assurance along with the Δ*G* calculation.

Figure 1 depicts the thermodynamic paths at the heart of SAFEP. The desired quantity 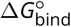 (red, left column) is equal to the sum of the steps in the SAFEP method (black, right column),

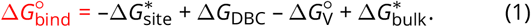

This equation forms the basis for the steps that follow in this tutorial.

**Figure 1.**
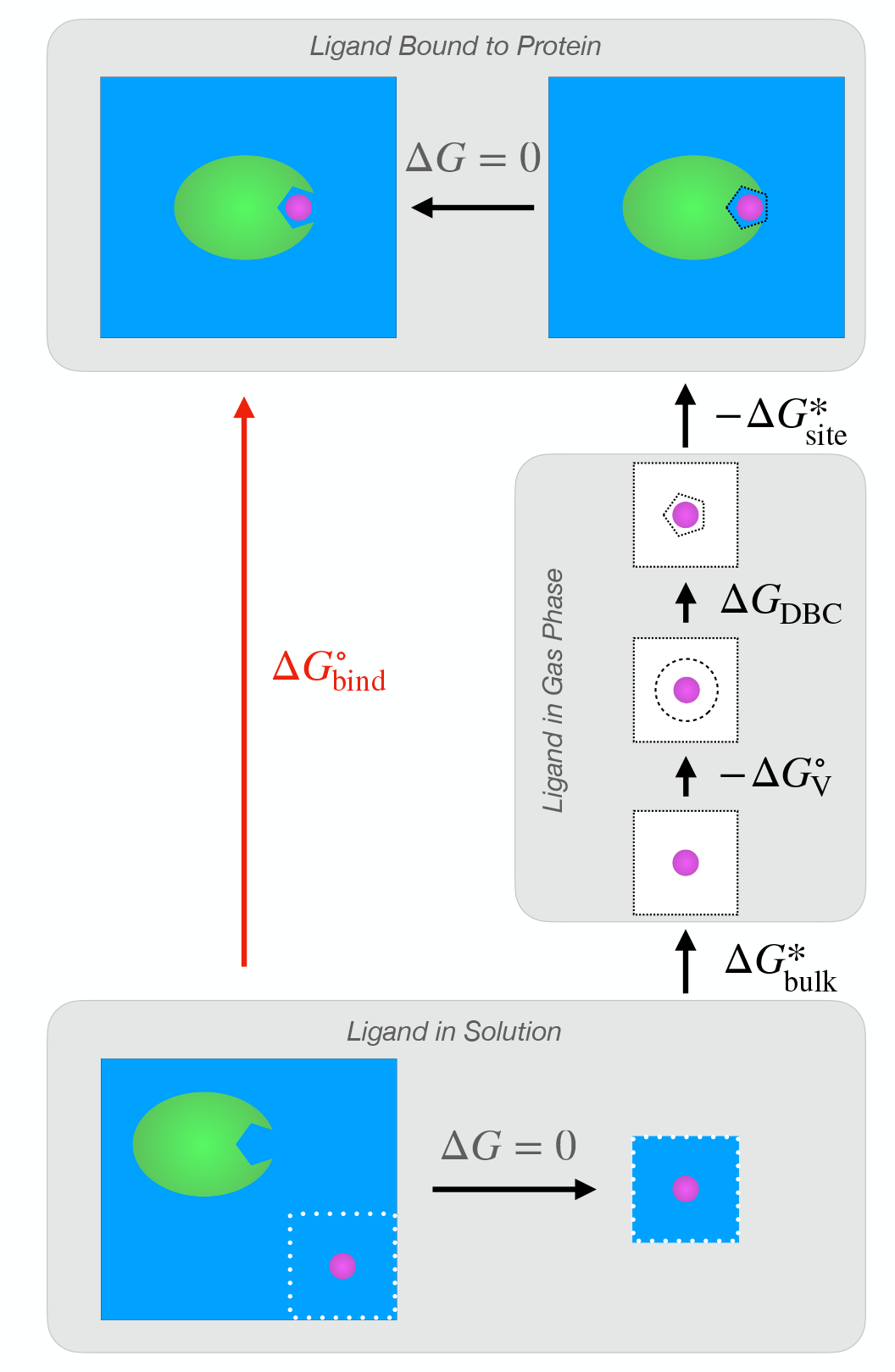
The SAFEP thermodynamic cycle. Computing the ABFE of a ligand bound to a protein 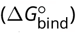 is the ultimate goal. This is found by computing the free energies of several, smaller perturbations: 1) decoupling the unbound ligand from the condensed phase to the gas phase under no restraints 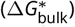 enforcing a restraint scheme (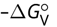 and Δ*G*_DBC_); and 3) coupling the ligand from the gas phase to its bound pose in the condensed phase 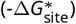 under the Distance-to-Bound Configuration (DBC) restraint. The free energy contribution of the volumetric restraint 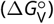 is calculated analytically, while the other three contributions are calculated via simulation. The free energy of the top horizontal leg vanishes in SAFEP due to the design of the DBC restraint. See Appendix C for more details.

### 1.1 Scope

The following steps will walk the user through the calculation of an Absolute Binding Free Energy (ABFE) using a computationally affordable example (phenol bound to a mutant lysozyme), but we have written these steps to be straightforward to generalize to other systems. These exact steps have been tested thoroughly for this particular system. More detailed discussions of each step can be found in Appendices A-E. To facilitate the generalization of the method to other systems, we have provided additional troubleshooting advice in Appendix F, as well as example files for each step. These appendix entries are hyperlinked and referenced throughout the body of the tutorial.

### 1.2 Prerequisites

#### 1.2.1 Background knowledge

We assume intermediate experience with running classical MD (any software) and beginner experience with NAMD 2.14 or later. If this is not the case, please see the NAMD Tutorial [15]. The latter portions that involve analysis are less important for this tutorial. Useful but not required material on alchemical free energy perturbations can be found in “*Insilico* alchemy: A tutorial for alchemical free-energy perturbation calculations with NAMD” [7]. Finally, basic knowledge of VMD and Python will be required for data analysis and visualization.

#### 1.2.2 Software requirements

1. NAMD2 version 2.14 or later, or NAMD3 version 3.0b3 or later. The NAMD3 series enables GPU-accelerated alchemical simulations. These instructions contain the command “namd2”: replace with “namd3” if applicable.
2. VMD 1.9.4.a57 or later. Slightly older versions of VMD may be used but will require a manual update of the Colvars Dashboard. See the Colvars Dashboard [README](https://github.com/Colvars/colvars/tree/master/vmd/cv_dashboard) for more information on getting the latest version.
3. Python 3.9.12 or later
4. Jupyter
5. safep Python package and its dependencies (see Procedure below for installation instructions)

NAMD will be used to perform simulations. GPU acceleration of restrained free energy perturbations are available in NAMD3 (with `CUDASOAintegrate off`) [16, 17]. System setup, trajectory visualization, and restraint definition will be carried out in VMD [18]. Data analysis and visualization will be handled by a Jupyter notebook with the above dependencies.

High-performance computing resources are recommended, but not required. Sample outputs are provided for each step for users with limited computing resources or time.

The Colvars Dashboard is the recommended tool for writing and editing Colvars config files. Advanced users can also edit the text files directly. Example files are provided for each step.

### 1.3 Process Overview

Within the scope of free energy perturbations, absolute free energies of binding are typically calculated by the double-decoupling method (DDM) [1, 3, 4]. In this method, pair interactions (non-bonded terms) between the ligand and the rest of the simulation box are gradually scaled to zero (decoupled) from both a bound state and an unbound, solvated state.

In order to maintain the ligand in its bound state, most current approaches introduce a series of rotational and translational restraints on the ligand, each of which requires parameterization and an additional Δ*G* correction. In contrast, SAFEP uses just one restraint: a flat-well on the “Distance-to-Bound Configuration” (DBC). This reduces both the number of parameters to be optimized and the number of simulations to be performed compared to layered restraints (See Figure 2 or Appendix C for details).

**Figure 2.**
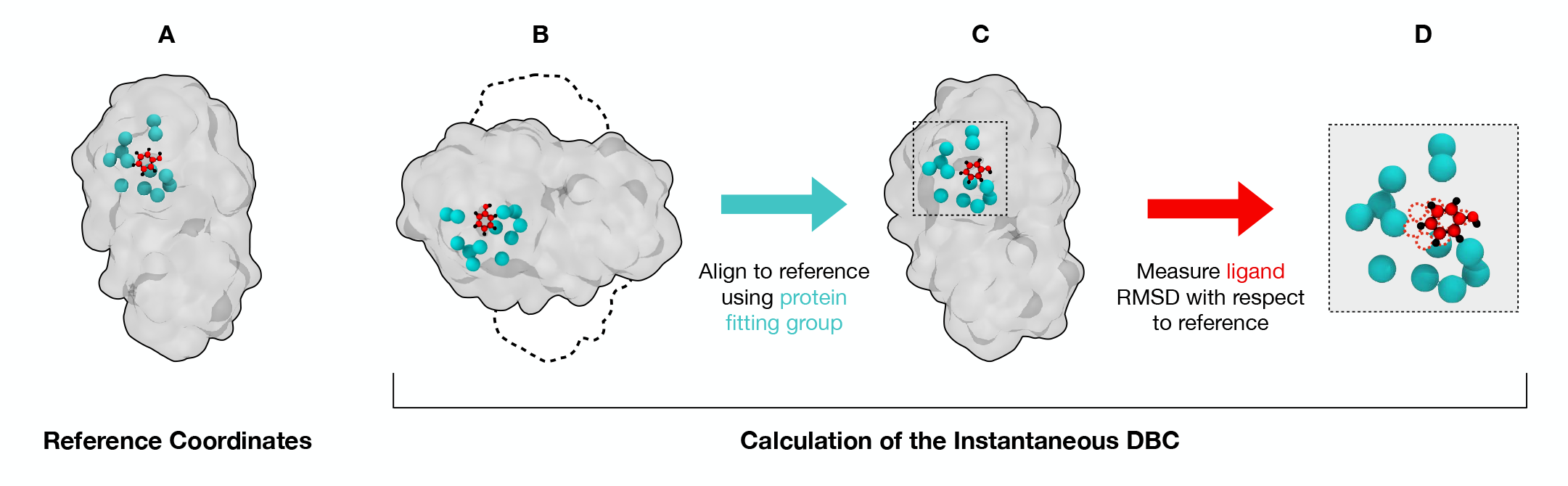
The Distance-to-Bound-Configuration (DBC) coordinate. The DBC coordinate is used as a bias to prevent ligand dissociation during uncoupling. The user specifies a subset of protein atoms as the fitting group (teal) and a subset of ligand atoms (red); also shown are the protein surface (gray) and remaining ligand atoms (black). A) User-specified reference coordinates for both protein and ligand. B) During simulation, both protein and ligand will drift from the reference coordinates (black dashed outline). C) In order to remove rotational and translational diffusion of the protein from the DBC calculation, Colvars aligns the system to the reference coordinates using only the protein fitting group atoms. D) The DBC is the RMSD of the user-specified ligand atoms (solid) with respect to the reference coordinates (dashed).

The thermodynamic cycle used for absolute binding free energies in SAFEP is shown in Figure 1 while the unknown values (black arrows) can be calculated by the simulations outlined in Figure 3. More precisely, the thermodynamic cycle (Fig 1) and the corresponding simulations (Fig 3) are broken into three main steps involving three simulation systems: 1) the ligand bound to the protein, 2) the ligand in the gas phase, and 3) the ligand in the bulk. The order of computations is unimportant so long as the endpoints are defined consistently (e.g. the same temperature is used throughout and restraints are used consistently). For the sake of clarity, we have arranged the process linearly: Steps A and B are concerned with calculating 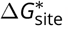, step C addresses the free energy of the DBC (Δ*G*_DBC_), step D measures 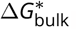, and step E calculates an analytical correction 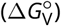 and combines all the preceding terms into the overall 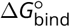 using Equation 1.

**Figure 3.**
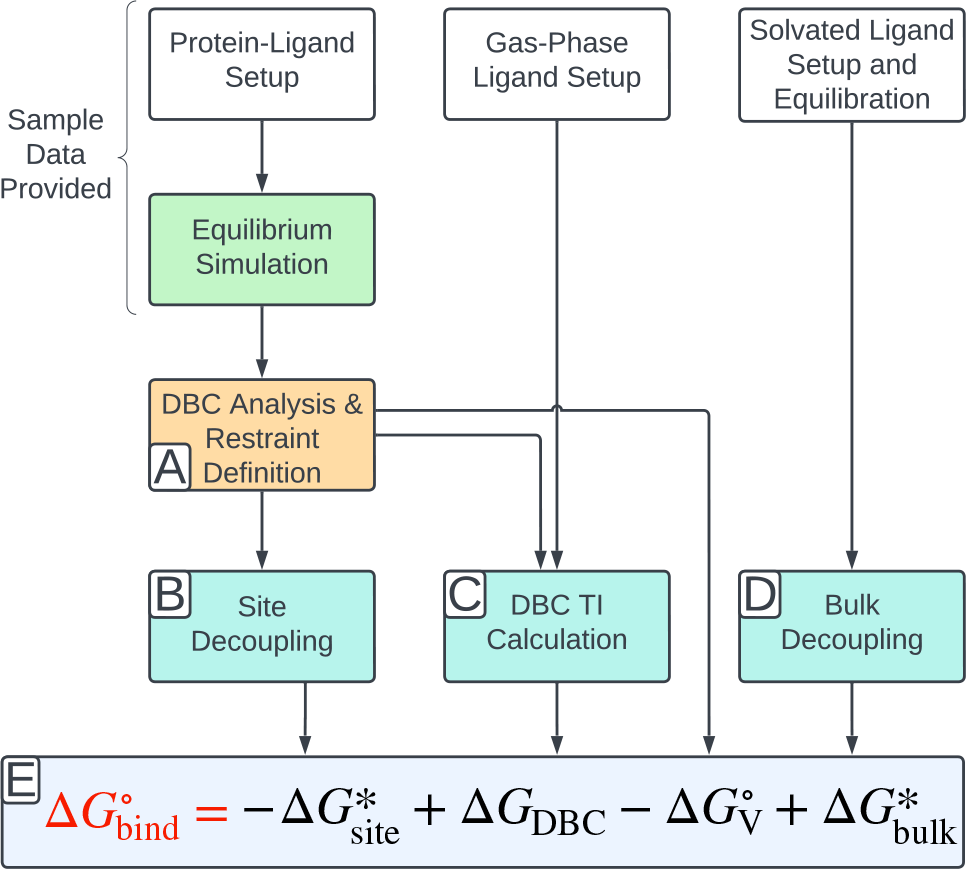
The overall SAFEP workflow. From top to bottom and left to right: 1) the ligand must be setup (as for classical MD) in each of the three states (bound, solvated, gas phase) and minimally relaxed (white boxes); 2) a longer, unbiased simulation of the ligand-protein complex is necessary to sample the bound state (green box) which is used to determine the distribution of the DBC (orange box, Step A); 3) two FEP calculations and a Thermodynamic Integration (TI) calculation are carried out (blue boxes, Step B, Step C, and Step D); and 4) the resulting values are combined to get the standard free energy of binding (gray box; Step E). Note: some simulations can be run simultaneously.

**Figure 4.**
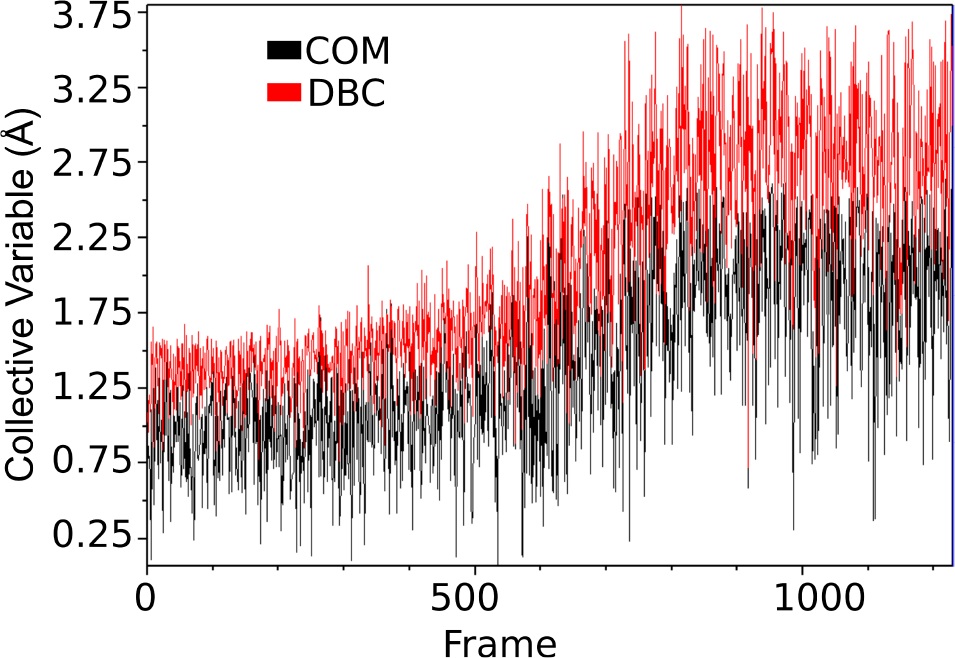
Screenshot of Colvars Dashboard output showing COM (red) and DBC (black) collective variables as a function of frame number in the decoupling trajectory from subsection 4.b in step C.

## 2 Protocol

The following steps demonstrate the SAFEP protocol applied to a computationally affordable example: calculating the binding affinity of phenol to a lysozyme mutant. For more details on the rationale behind this choice, see Appendix A.

**Procedure:**

1. **Clone the SAFEP_tutorial repository to your local environment** Tip: use `--depth 1` to avoid downloading the entire commit history.

~~~
git clone https://github.com/jhenin/SAFEP_tutorial.git--depth 1
~~~
2. Navigate to the cloned repo
This is the starting path for all command line prompts in this tutorial.

~~~
cd SAFEP_tutorial
~~~
3. Install the SAFEP package by running:

~~~
pip install git+https://github.com/BranniganLab/safep.git
~~~
4. **Tips:**
  - All run.namd files contain a line that reads `set useSampleFiles 0`. To use the sample data provided, set the value to 1. Otherwise, NAMD will use your inputs (provided they are named exactly as described in this document).
  - If you run VMD and your simulations on different computers, then you will need to manually edit paths later when you are running simulations.
  - Some simulations will take several days on a single core. To use 4 cores in parallel we have included the +p4 argument in the commands for the longer NAMD runs. This number may need to be optimized for your particular computing resources.
  - If you are using namd3 with CUDA (GPU) acceleration, you should consult the NAMD User Guide [13] for optimal settings. As of NAMD3b3, CUDAOSOAIntegrate should be left OFF, and “+idlepoll” should be included in the run command.
  - Common settings used by multiple simulations are in *common/common_config*.*namd*, which is sourced by the individual configuration files. This simplifies the individual configuration files and ensures consistency between calculations, which is a critical part of any free energy method.
5. **Move on to Step A**

### Step A: Sample the bound state and define the corresponding restraint

Alchemical decoupling removes the interactions that stabilize the occupied ensemble. Consequently, during decoupling the ligand may spontaneously diffuse into the bulk. Therefore we need to impose an external restraint to force the ligand to occupy the bound state throughout decoupling. With SAFEP we apply a single restraint on the Distance-to-Bound Configuration (DBC) collective variable as illustrated in Fig. 2. This restraint is straightforward to define and relatively insensitive to small differences in parameters [11]. Its sole correction factor is calculated via Thermodynamic Integration later in this tutorial. For more details about the DBC restraint see Appendix C and [11]. For a discussion of the merits of accounting for symmetry when computing the DBC of a small, symmetric ligand see Appendix C.2.2 and Ref. 19.

#### Required Input

- Structure file: *common/structures/phenol_lysozyme*.*psf*
- Coordinate file: *common/structures/phenol_lysozyme*.*pdb*
- Equilibrium trajectory: *stepA_create_DBC/inputs/unbiased-sample*.*dcd*

#### Essential Output

- DBC restraint parameters: *stepA_alchemy_site/outputs/DBC_restraint*.*colvars*

#### Procedure

1. *Note: We have completed this step for you in the specific case of this tutorial*. **Run standard MD of the occupied state**. This simulation should be long enough (∼50 ns) for the ligand to explore the configuration space of the bound state. See Appendix A for more details.
2. **Define the Distance-to-Bound Configuration (DBC) Coordinate**
  a. **Open the Colvars Dashboard in VMD:**
    - Open VMD.
    - Load the psf, pdb, and dcd files listed above under “Required Input.” You may choose to load your own dcd if you completed Step 1.
    - From VMD’s main window options select: *Extensions*→*Analysis*→*Colvars Dashboard*
  b. **Create a DBC colvar:**
    - Click 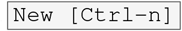 to start editing a new collective variable.
    - Delete all sample text shown in the editor on the right-hand side.
    - Open the *Templates*→*colvar templates* drop-down list and select `DBC (ligand RMSD)` to populate the editor with a template that now needs to be edited.
  c. **Define the atom selection for the ligand atoms: (Fig. 2, red)**
    - Delete `atomNumbers 1 2 3 4` from the `atoms` block and leave your cursor on the now-empty line.
    - Select the left panel text box *Editing helpers*→*Atoms from selection text* and enter `resname PHEN and noh`.
    - Press Enter or click 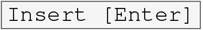 to insert the new selection into the configuration text at your cursor.
  d. **Identify equivalent, symmetric atoms:**
    - In the rmsd block, add `atomPermutation 1 5 3 9 7 11 12` to the line above the atoms keyword. This indicates equivalence between ligand atoms listed in `atomNumbers`. That is, (5 and 3) and (9 and 7) are interchangeable. See the Colvars User Guide and Appendix C.2.2 for more details on symmetric DBC and `atomPermutation`.
  e. **Define the atom selection for the binding site atoms: (Fig. 2, teal)**
    - Delete `atomNumbers 6 7 8 9` within the `fittingGroup` block and leave your cursor on the now-empty line.
    - Select the left panel text box *Editing helpers*→*Atoms from selection text* and enter `alpha and same residue as within 6 of resname PHEN`.
    - Press Enter or click 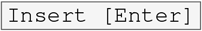 to insert the new selection into the configuration text at your cursor.
    - **VMD versions 1.9.4a57 and older only** Colvars has a functionality called “auto-update selections.” Please turn it off by deleting the comment line that begins with “auto-updating” in all Colvars config files. It is off by default in newer CV Dashboard versions.
  f. **Set the reference positions for the RMSD and alignment calculations:**
    - In the initial `RMSD` block, before the `atoms` block, delete `refpositionsfile reference.pdb # PDB or XYZ file` (the first highlighted line in the figure below). Leave your cursor on that line.
    - In the left panel under *Editing helpers*, select the radio button ° `refPositionsFile` and click 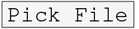.
    - Select the *phenol_lysozyme*.*pdb* file you used as input for this section. This will insert a line in the dashboard text editor that indicates the file that will be used for the DBC reference coordinates.
    - Copy the line just inserted and replace the `refpositionsfile` line at the bottom of the `atoms` block (the second highlighted line in the figure below). This sets the same PDB file to be used for aligning to the protein frame-of-reference.
    - For NAMD builds older than October 31, 2022: change “centerToReference” and “rotateToReference” to “centerReference” and “rotateReference” respectively.
    - The colvar config editor should now look like the screenshot below with your file’s path in place of the two highlighted lines. 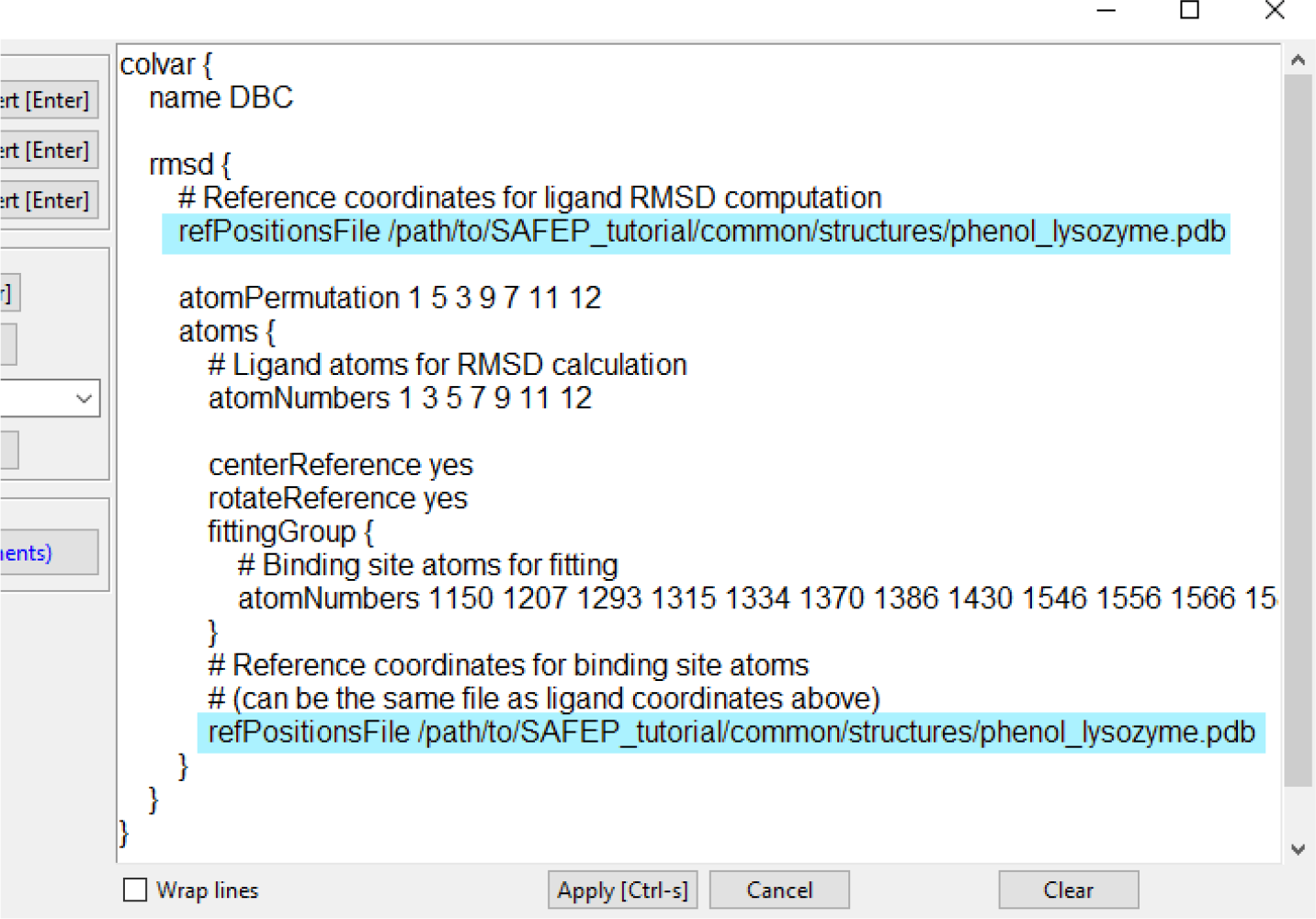
  g. **Save your edits:**
    - Click the 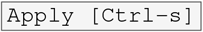 button.
3. **Impose a restraint based on the DBC coordinate**
  a. **Determine the upper wall of the DBC restraint:**
    - In the *Plots and real-time visualizations* panel of the dashboard, click 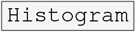. If you don’t see such a button, you need to upgrade your VMD installation. See Software requirements for more details.
    - From the histogram, estimate the 95th percentile of the bound state’s DBC coordinate. Use the cumulative distribution line graph as a guide. The value doesn’t need to be precise. We selected 1.5 Å. See Appendix C.2.2 for more details.
    - Write this value down; you will need it in the next step.
  b. **Impose a flat-bottom harmonic potential:**
    - Open the *Biases* tab on the Colvars Dashboard and click 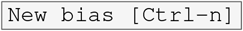 to create a new biasing potential.
    - Delete the default text.
    - From the *bias templates:* drop-down menu select `harmonicWalls` and click 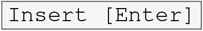.
    - Modify the bias to match the following parameters: The force constant in this case is in units of kcal/(mol·Å^2^). The strength of restraint should be neither so great that it causes instabilities nor so weak that it fails to cleanly separate the bound and unbound ensembles.
  c. **Save your edits:** Click the 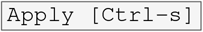 button.
4. **Save the Colvars configuration to a file**
  a. **Click** 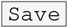 **at the top of the dashboard**
  b. **Save your file to** *stepA_create_DBC/outputs/DBC_restraint*.*colvars* Note that if you choose to use a different file name or path you will need to update the files in the next step with the new name.

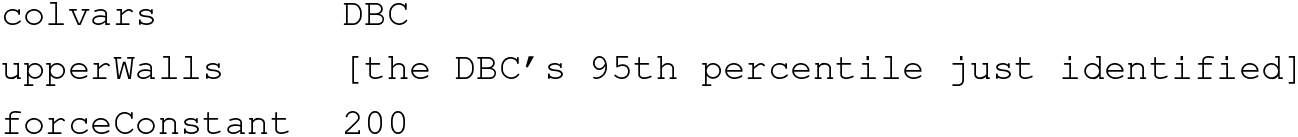

### Step B: Decouple phenol from the protein via FEP

In this section we will calculate 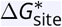 by decoupling the ligand from the protein binding site (and all other contents of the simulation box) using alchemical FEP. We will maintain the ligand in the bound configuration relative to the protein by restraining the DBC coordinate as defined in the previous step.

#### Required Input

- Structure file: *common/structures/phenol_lysozyme*.*psf*
- Coordinate file: *common/structures/phenol_lysozyme*.*pdb*
- DBC restraint parameters: *stepA_create_DBC/outputs/DBC_restraint*.*colvars*
- NAMD configuration file: *stepB_alchemy_site/inputs/run*.*namd*

#### Essential Output

- FEP configuration file: *stepB_alchemy_site/outputs/alchemy_site*.*pdb*
- FEP trajectory file: *stepB_alchemy_site/outputs/alchemy_site*.*dcd*
- FEP output file: *stepB_alchemy_site/outputs/alchemy_site*.*fepout*

#### Procedure

1. **Specify which atoms will be decoupled using the pdb beta field**
  a. **Open VMD and load the psf and pdb files specified in “Required Input.”**
  b. **Set and write beta values:**
    - Open the Tk Console from the Extensions menu.
    - Ensure that your Tk Console is in the correct directory:

~~~
cd stepB_alchemy_site/outputs
~~~
    - Set the beta value of all atoms to 0:

~~~
[atomselect top all] set beta 0
~~~
    - Set the beta values of the ligand atoms to -1 for decoupling:

~~~
[atomselect top “resname PHEN”] set beta -1
~~~
    - Save as a pdb file:

~~~
[atomselect top all] writepdb alchemy_site.pdb
~~~
2. **Perform the FEP simulation** We have provided a configuration file for this FEP run: *stepB_alchemy_site/inputs/run*.*namd*. See the in-line comments in that file and Appendix B for a detailed description of the settings relevant to running FEP in namd.
  a. **Run the decoupling FEP**: Enter the following in your terminal window:

~~~
cd stepB_alchemy_site/inputs/
namd2 +p4 run.namd > ../outputs/alchemy_site.log 2> ../outputs/alchemy_site.err
~~~
  b. **[Optional] Start Step C:** If you have access to more compute resources, you can continue on to Step C while this FEP calculation is running. **Don’t forget to return to analyze these data once the simulation is complete**.
3. **Analyze the trajectory**
  a. **Visually inspect the trajectory in VMD:**
    - Open VMD.
    - Load the .psf (*common/structures/phenol_lysozyme*.*psf*) and .dcd file(s) from the outputs of stepB.
    - Ensure that the ligand remains in a bound-like configuration for the duration of the simulation.
  b. **Measure the restraint energy:**
    - Open the Colvars Dashboard.
    - Click 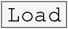 and import your DBC restraint file (*DBC_restraint*.*colvars*).
    - Open the *biases* tab, select the DBC restraint, and click 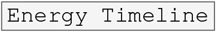.
    - The restraint energy should remain near zero for several nanoseconds, then increase and reach a maximum in the second half of the simulation (when the ligand is fully decoupled). If this is not the case, see Appendix F.
  c. **Calculate** 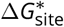 **in the Jupyter Notebook:**
    - Navigate back to the tutorial root directory.
    - Begin a Jupyter session and open the notebook titled *SAFEP_Tutorial_Notebook*.*ipynb*.
    - Follow the in-notebook prompts to parse your new fepout file (*stepB_alchemy_site/output/AFEP2-02*.*fepout*). By default, we use the sample output. Be sure to update the paths as indicated in the notebook: 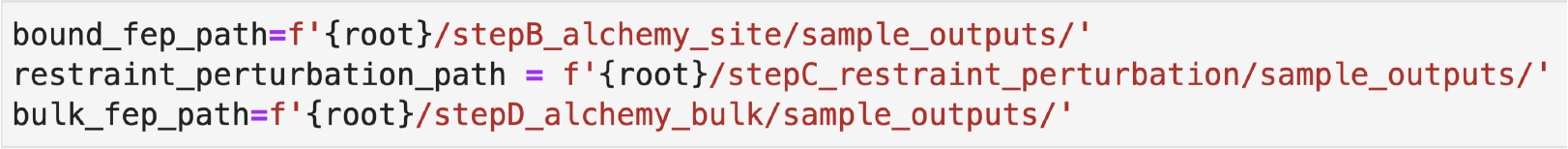
    - Compare your outputs to the sample outputs found in Appendix B.3.
    - Caution: the errors reported are based on the PyMBAR BAR estimator and are subject to the corresponding assumptions and caveats [20].

#### Step C: Compute the DBC restraint free energy correction

We designed the DBC restraint so that it doesn’t do any significant work in the fully coupled system. However it does reduce the entropy of the fully decoupled ligand, which would otherwise be exploring an “empty” simulation box. We need to calculate the corresponding free energy cost so we can correct for it. In this section we will use thermodynamic integration (TI) to calculate Δ*G*_DBC_: the free energy difference between a gas-phase ligand under DBC restraints *v*s a (spherical) volumetric restraint. For more details see Appendix D.

##### Required Input

- Structure file: *common/structures/phenol_gas_phase*.*psf*
- Coordinate file: *common/structures/phenol_gas_phase*.*pdb*
- NAMD configuration file: *stepC_restraint_perturbation/inputs/run*.*namd*

##### Essential Output

- Colvars configuration file: *stepC_restraint_perturbation/outputs/DBC_Restraint_RFEP*.*colvars*
- FEP trajectory file: *stepC_restraint_perturbation/outputs/RFEP*.*dcd*
- Colvars output file: *stepC_restraint_perturbation/outputs/RFEP*.*colvars*.*traj*

**Procedure:**

1. **Define the collective variables**
  a. **Open VMD and load the input files:**
    - Open VMD.
    - Open the Tk Console.
    - Open the Colvars Dashboard.
    - **[Optional]** Extract the phenol from the phenol-lysozyme complex by running the following in the tkConsole. **Note: We have completed this step for you. The sample files can be found in *common/structures***.

~~~
%cd common/structures
%mol load psf phenol_lysozyme.psf pdb phenol_lysozyme.pdb
%set ligand [atomselect top “resname PHEN”]
%cd ../../stepC_restraint_perturbation/outputs
%$ligand writepsf phenol_gas_phase.psf
%$ligand writepdb phenol_gas_phase.pdb
~~~
    - Load *phenol_gas_phase*.*psf* and *phenol_gas_phase*.*pdb*
  b. **Define the gas-phase spherical coordinate:**
    - In the Colvars Dashboard, click 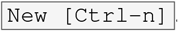.
    - In the second line of the editor, replace the default name `myColvar` with `COM`.
    - Delete `atomNumbers 1 2` and leave your cursor on that line.
    - Using the *Atoms from selection text:* tool in the left panel, enter `resname PHEN and noh` and click 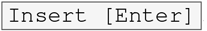.
    - Get the geometric center of the heavy atoms by the following in the Tk Console:

~~~
measure center [atomselect top “resname PHEN and noh”]
~~~
    - Set the atoms of group 2 to dummyAtom (x0, y0, z0) where x0, y0, and z0 are the coordinates of the geometric center of the ligand you just retrieved in the previous step. Your editor should look similar to the figure below. Note the inclusion of commas in the `dummyAtom` statement. 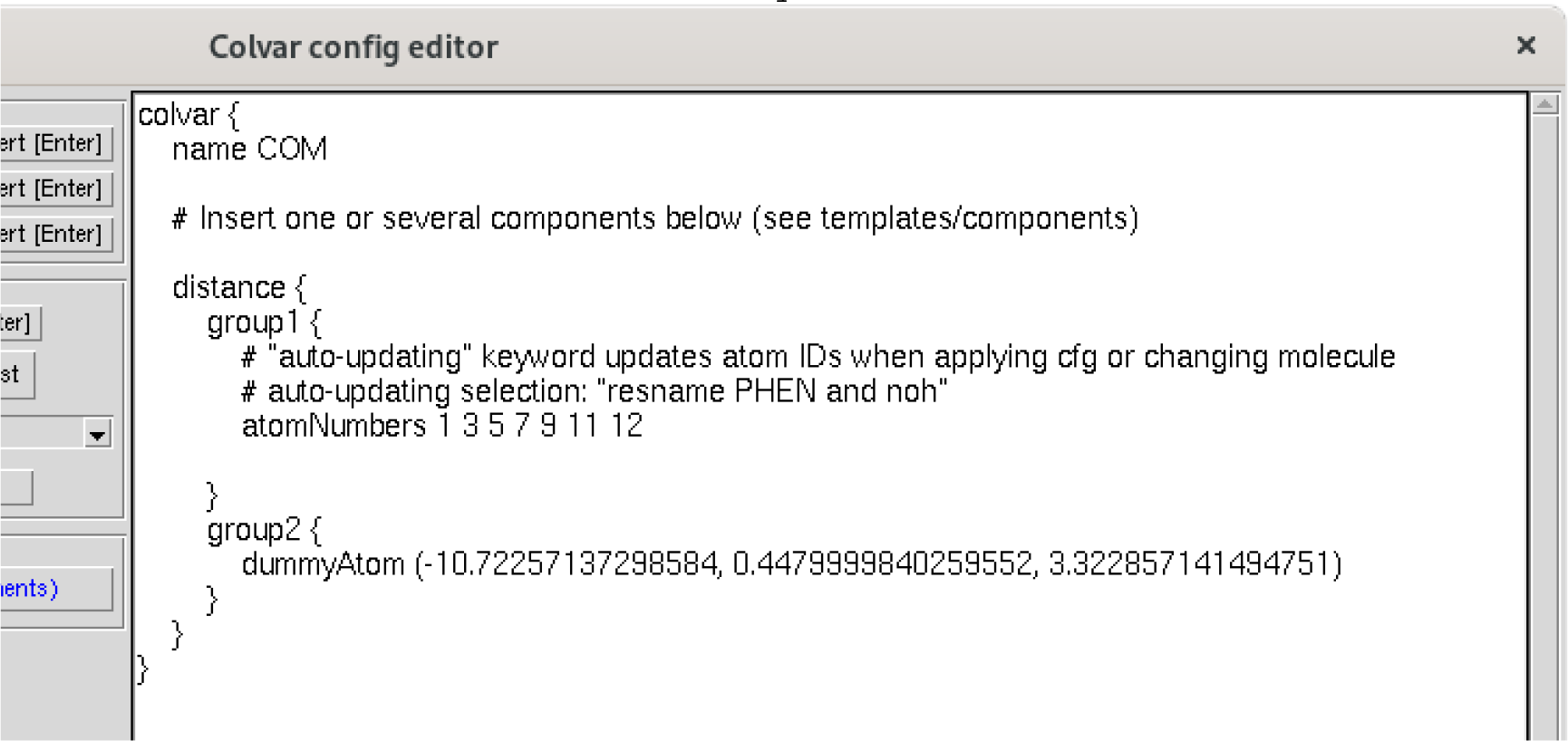
    - Save and close the colvar editor by clicking 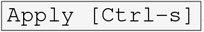.
  c. **Define the gas-phase DBC coordinate:**
    - Click 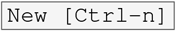again.
    - In the second line of the editor, replace the default name `myColvar` with `DBC`.
    - Delete the default distance component `distance{…}` and leave your cursor on that line.
    - From the *component templates* dropdown menu select `rmsd` and click 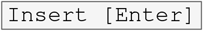.
    - As before, add `atomPermutation 1 5 3 9 7 11 12` to the rmsd block to define the ligand symmetry.
    - Delete `atomNumbers 1 2 3` and leave your cursor on that line.
    - In the field labeled *Atoms from selection text:* enter `resname PHEN and noh` and click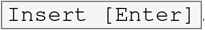.
    - Add `rotateReference off` and `centerReference off` to the `atoms` block.
    - Replace the default refPositionsFile @ line using the ° `refPositionsFile` radio button and the 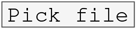 button to select *phenol_gas_phase*.*pdb*.
    - Save and close the colvar editor by clicking 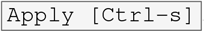.
2. **Define the restraints**
  a. **Create the spherical restraint:**
    - In the *biases* tab of the Colvars Dashboard, click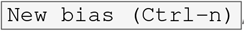, and delete the default text.
    - From the *bias templates* dropdown menu, select `harmonic walls`, and press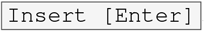.
    - Recall the upperWalls value you used for the DBC restraint in subsection 3.b from Step A. You will need this value in this and the next step.
    - Modify the bias to match the following parameters (see Appendix D):
    - Save and close the bias editor by clicking 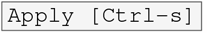.
  b. **Save the config file:**
    - Click the 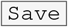button on the Colvars Dashboard.
    - Save the file as *stepC_restraint_perturbation/outputs/DBC_restraint_RFEP*.*colvars*
  c. **Create a DBC restraint that gradually releases:**
    - We will use the provided setTI Tcl procedure.
    - Open stepC_restraint_perturbation/inputs/run.namd in a text editor
    - Find the block labeled “COLVARS”
    - Edit the input variables to match the following
3. **Run the TI simulation**
  a. **Enter the following in your terminal:**

~~~
cd stepC_restraint_perturbation/inputs
namd2 +p1 run.namd > ../outputs/DBC_FreeEnergy.log 2> ../outputs/DBC_FreeEnergy.err
~~~
  b. **[Optional] Start Step D:** If you have access to more computing resources, you can continue on to Step D while the TI calculation is running. **Don’t forget to return to analyze these data once the simulation is complete**.
4. **Analyze the output** If any of these checks fails, check the Troubleshooting section of the Appendices (Appendix F).
  a. **Visually inspect the trajectory in VMD:**
    - Open VMD.
    - Load the .psf, .pdb, and .dcd files associated with this tutorial step.
    - The ligand should initially fluctuate roughly in place at the start and gradually explore the COM restraint space as the DBC restraint is released.
  b. **Check the collective variable trajectories:**
    - Open the Colvars Dashboard
    - Click 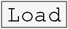 and open the Colvars configuration file *DBC_restraint_RFEP*.*colvars*
    - Select both the COM and DBC restraints
    - Click 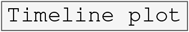
    - Both coordinates should start low and gradually increase. The COM restraint should plateau near the value of the restraint’s upperWalls as shown in the figure below:
  c. **Calculate** 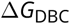**in the Jupyter Notebook:**
    - Open the Jupyter Notebook as in subsection 3.c from step B. Update the paths in “User Settings” as needed. Remember, sample data will be used by default.
    - Run the first several cells at least until the first FEP analysis section.
    - Run the first cell in the section titled “Process the DBC TI calculation” and make sure the DBC and COM walls are correct.
    - Run all the remaining cells in that section. The output will include Δ*G*_DBC_ as well as an error estimate. Sample results and more explanation can be found in Appendix D.

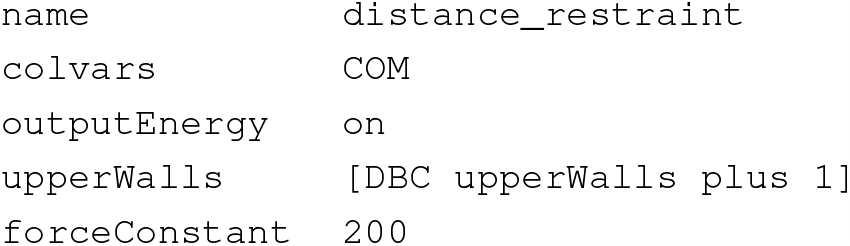

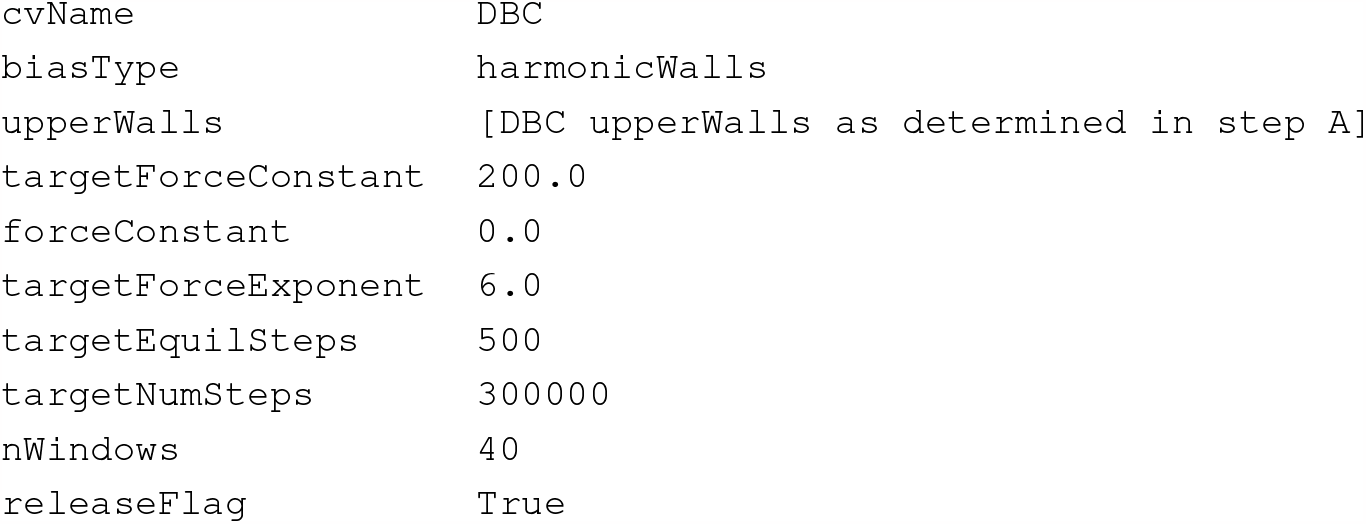

#### Step D: Decouple phenol from bulk solvent

You have completed one alchemical FEP calculation already, but double-decoupling methods require two such calculations to close the thermodynamic cycle. We need to know the free energy of transferring the ligand from the binding site into vacuum, *and* from vacuum into the bulk. In this section we will calculate the latter term, 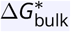, by decoupling the ligand from the bulk solution.

##### Required Input

- Structure file: *common/structures/phenol_water*.*psf*
- Coordinate file: *common/structures/phenol_water*.*pdb*
- NAMD configuration file: *stepD_alchemy_bulk/inputs/run*.*namd*

##### Essential Output

- FEP configuration file: *stepD_alchemy_bulk/outputs/alchemy_bulk*.*pdb*
- FEP trajectory file: *stepD_alchemy_bulk/outputs/alchemy_bulk*.*dcd*
- FEP output file: *stepD_alchemy_bulk/outputs/alchemy_bulk*.*fepout*

**Procedure:**

1. *Note: We have completed this step for you in the specific case of this tutorial*. **Prepare the ligand as you would for a traditional simulation**. Use a sufficiently large box size; very small boxes are more prone to instabilities and self-interactions which will lead to artifacts in the final free energy estimate.
2. **Specify which atoms will be decoupled using the pdb beta field**
  a. **Open VMD and load the psf and pdb files specified in “Required Input.”**
  b. **Set and write beta values:**
    - Open the Tk Console
    - Ensure that your Tk Console is in the correct directory:

~~~
cd stepD_alchemy_bulk/outputs
~~~
    - Set the beta value of all atoms to 0:

~~~
[atomselect top all] set beta 0
~~~
    - Set the beta values of the ligand atoms to -1 for decoupling:

~~~
[atomselect top “resname PHEN”] set beta -1
~~~
    - Save as a pdb file:

~~~
[atomselect top all] writepdb alchemy_bulk.pdb
~~~
3. **Run the ligand decoupling simulation in bulk solvent**

~~~
cd stepD_alchemy_bulk/inputs
namd2 +p4 run.namd > ../outputs/alchemy_bulk.log 2> ../outputs/alchemy_bulk.err
~~~
4. **Analyze the output**
  a. **Visually inspect the trajectory in VMD:**
    - Open VMD.
    - Load the .psf, .pdb, and .dcd files associated with your simulation.
    - The ligand should diffuse normally at the start of the simulation but behave more and more like a gas-phase molecule.
  b. **Calculate** 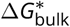 **in the Jupyter Notebook**
    - Open the Jupyter Notebook as in subsection 3.c from Step B.
    - Confirm that bulk_fep_path points to your files
    - Parse the *fepout* file by running all the cells in the Jupyter notebook section titled “Decoupling from Solvent.”
  c. Caution: as noted above, the errors reported are based on the PyMBAR BAR estimator and are subject to the corresponding assumptions and caveats [20].

#### Step E: Calculate corrections and combine quantities

We will now calculate 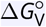 analytically. With this final piece of information, we can calculate the dissociation constant and estimate a titration curve based on the probability of occupancy assuming a two-state system: 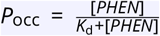 where *K*_d_ is the dissociation constant given by 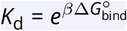. For additional information see Appendix E.

##### Required Input

- Site FEP data: *stepB_alchemy_site/outputs/alchemy_site*.*fepout*
- Restraint perturbation data (RFEP/TI): *stepC_restraint_perturbation/outputs/RFEP*.*colvars*.*traj*
- Bulk FEP data: *stepD_alchemy_bulk/outputs/alchemy_bulk*.*fepout*

##### Essential Output

- 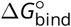
- *titration_curve*.*pdf*

##### Procedure

1. Complete any unfinished analyses in previous steps (i.e. steps B.3, C.4, and D.4). Some issues are only obvious when visualizing the trajectories. Don’t skip those steps!
2. Open the Jupyter notebook and navigate to the section labeled “Volumetric Restraint Contribution.”
3. Run the section to calculate the volumetric free energy contribution. See Appendix C.3 for a more detailed explanation. **Note:** At this point you will either need to have completed all simulations or use the sample data provided. To use the sample data, change the path variables (`bound_fep_path, restraint_perturbation_path, and bulk_fep_path`) to use the files in their respective .*/sample_outputs* directories.
4. Calculate the overall 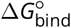 and compute a titration curve by running the cells in the section “Binding Free Energy.”
5. Compare your final 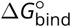 to the literature value, reported as -5.44 kcal/mol [21].
6. Compare your titration curve to Figure 5 below.

**Figure 5.**
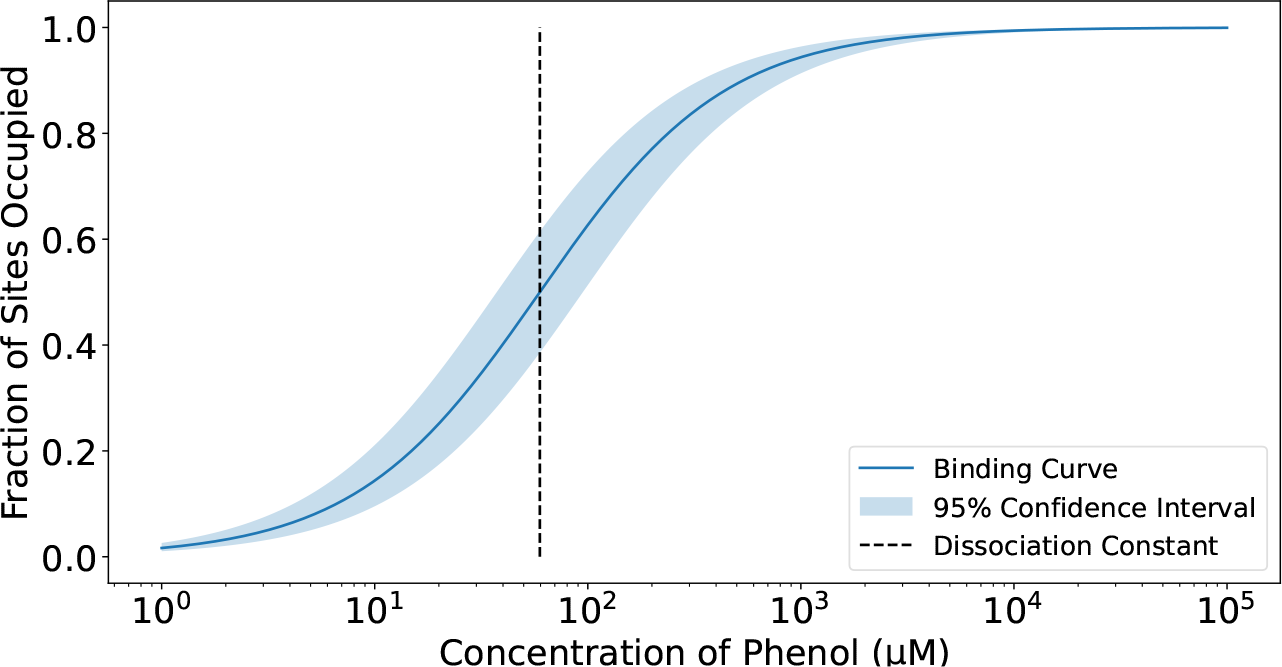
An example titration curve. generated using Equation 23. The 95% confidence interval represents *±*1.96 *∗* SEM of Δ*G*_bind_.

### 3 What’s next?

Congratulations! You have completed all the essential steps of the SAFEP protocol. You can now apply the protocol above to your own receptor-ligand binding problem, applying minor changes as needed. Of course, the outcome of your SAFEP calculation will be only as realistic as your model and force field. As you proceed, it is critical to visualize the simulations, track the evolution of the DBC, and evaluate the convergence of your own calculations. For a more in-depth analysis of the alchemical calculations, see the general-purpose notebooks from the SAFEP repository at https://github.com/BranniganLab/safep. Additional information and a (non-exhaustive) troubleshooting guide can be found in the appendices below. If you have trouble with this tutorial or the SAFEP package, please contact us either directly or by opening an issue on the corresponding GitHub page. We anticipate that future updates to this tutorial will incorporate lessons from user feedback and our own experience, including new examples requiring changes in the protocol.

**Table 1.** Symbols used in this tutorial.

| Symbol | Definition |
| --- | --- |
| $d$ | Distance-to-Bound-Configuration (DBC) coordinate |
| $d_i$ | $d$ for ligand $i$ |
| $\Delta G_\lambda$ | The change in (Gibbs) free energy for a particular $\lambda$ window |
| $\Delta G_{\text{bind}}^\circ$ | Standard free energy of binding |
| $\Delta G_{\text{bulk}}^*$ | Free energy of decoupling the ligand from the bulk solution |
| $\Delta G_{\text{DBC}}$ | Free energy of imposing the DBC restraint on the ligand from a COM restraint |
| $\Delta G_{\text{site}}^*$ | Free energy of decoupling the ligand from the binding site |
| $\Delta G_V^\circ$ | Free energy of releasing the ligand to the standard state from the center of mass restraint |
| $K_d$ | The dissociation constant for the ligand-protein complex |
| $k$ | A force constant for a restraint |
| $k_\lambda, k_0$, and $k_1$ | A force constant that is a function of $\lambda$, when $\lambda = 0$, and $\lambda = 1$, respectively |
| $L$ and $L'$ | Arbitrary ligand concentrations |
| $m$ | Number of non-decoupled ligands in a bulk system |
| $n$ | Number of (TI) $\lambda$ windows |
| $P_{\text{occ}}$ | The probability of the binding site being occupied at a particular ligand concentration, defined in Eq. 21 |
| $r$ | Distance of ligand center of mass (COM) from a reference |
| $r_R$ | Radius of a flat-well COM restraint |
| $U$ | Potential energy |
| $U_{\text{FW}}$ | Potential energy of a flat-well bias |
| $V$ | Volume of the bulk system |
| $V_R$ | Volume of a sphere with radius $r_R$ |
| $Z_{\text{occ}}$ and $Z_{\text{unocc}}$ | Partition functions for the occupied state and unoccupied state, respectively, defined in Eqs. 13 and 14 |
| $\alpha$ | An exponential smoothing parameter |
| $\beta$ | $(k_B T)^{-1}$ |
| $\delta_{\lambda,i}$ | The free energy discrepancy between forward and backward sampling of $\lambda$ window $i$, defined in Eq. 2 |
| $\Theta_i$ | DBC test function for a single ligand molecule in the system, defined in Eq. 11 |
| $\Theta_{\text{occ}}$ | Occupancy test function for a single binding site, defined in Eq. 12 |
| $\Delta \lambda_i$ | The width of the $i$th $\lambda$ window |
| $\lambda$ | A coupling parameter $\lambda \in \{0, 1\}$ |
| $\xi$ | An arbitrary collective variable |
| $\xi_{\text{max}}$ | Upper wall of a flat-well restraint on $\xi$ |
| $\sigma_{\Delta G}$ | The standard deviation of the total free energy, defined in Eq. 9 |
| $\sigma_i$ | The standard deviation of the free energy for a particular state $i$, defined in 10 |

## Appendix A System Selection and Setup

Lysozyme L99A/M102H (PDBid 4I7L) was chosen for several reasons. Lysozyme L99A/M102H is a small protein that binds a small, rigid molecule with reasonably high affinity which has already been measured experimentally. These properties make it well-suited as a model for prototyping and validating free energy calculation methods generally.

Because lysozyme is elongated, we save some computation time by using a narrower box. Rotations of the protein could cause self-interactions, which we avoid by imposing a soft harmonic restraint on its orientation, as defined in *common/protein_tilt*.*colvars*. The provided systems were prepared using CHARMM-GUI [22, 23] using a truncated lysozyme (PDBid 4I7L, residues 3 to 157) and solvated using default parameters (TIP3P water, 0.15 M NaCl). The production run uses largely default parameters and settings. The only notable exception is that WrapAll should be set to off. This is because wrapping across the PBC can cause unexpected results during analysis which can compromise the FEP and TI calculations.

## Appendix B Running FEP in NAMD

### Appendix B.1 Configuration Files

In addition to the configuration, forcefield, and structural files, running FEP in NAMD requires a particular pdb file (sometimes called a “fep file”) that contains flags that indicate which atoms are being coupled or decoupled. This is usually indicated in the beta column as ‘-1’ for decoupling or ‘1’ for coupling. All other atoms should have beta set to 0.

The configuration file also contains some additional options that are detailed in the NAMD user guide [13] and described briefly in the provided configuration files. While most of the settings should remain unchanged in a wide range of systems, there are a few exceptions.

**alchOutFreq** determines the number of steps between collecting FEP samples. It should be set to a multiple of `fullElectFreq` to ensure accurate energy estimates. Later versions of NAMD should resolve this issue automatically; see [Bug advisory and Workaround](https://www.ks.uiuc.edu/Research/namd/mailing_list/namd-l.2020-2021/1487.html). Additionally, the sampling frequency should be between 50 and 200 steps; sampling too frequently will result in bloated data sets of highly autocorrelated samples while sampling infrequently will result in too few samples to get a well-converged estimate of the change in free energy.

**alchVdwShiftCoeff** controls the strength of the softcore potential which is essential to prevent “end-point catastrophes” in which one or more Lennard-Jones potentials diverge to infinity near λ=0 or λ=1. In example inputs for this tutorial, we set it to 6 Å^2^, which is a safe value for a range of molecular systems. See [24] and [19] for more details.

**alchElecLambdaStart** is the value of λ for which discharge of the ligand is complete, while Lennard-Jones interactions begin weakening when λ equals **alchVdwLambdaEnd**. In this tutorial, we have set both parameters to 0.5, which separates the discharge phase of the calculation from the Lennard-Jones decoupling, and works well for many cases.

**alchEquilSteps** determines the time between starting a new λ value and beginning to sample the ensemble. Alchemlyb and PymBAR provide functions that will down-sample the data set using automated equilibrium detection schemes. We have found that automated equilibrium detection performs about as well as manually setting `alchEquilSteps` and autocorrelation is the bigger problem when trying to assess convergence. As a result we have set this to 1. See [20] for a more detailed discussion of equilibrium detection, autocorrelation, and their effects on free energy estimation. See Appendix B.3 or the provided Jupyter notebook for more information on how these are used in our analysis.

**deltaLambda** is passed as a parameter to the runFEP function and determines the width of the λ windows. Narrower windows will converge faster but will increase the total number of windows required to span λ = 0 to λ = 1. As a result, we need to empirically optimize the number and length of windows. See Appendix B.3 and Appendix F for more details on assessing and optimizing these parameters. The number and length of windows used here (∼ 40 ns total simulation time) are a good starting point, but very flexible ligands or dynamic binding sites may require much more sampling.

**IDWS** (interleaved double-wide sampling) tells NAMD to alternate between sampling the forward and reverse λ directions (via the runFEP function, which is passed the `lchLambdaIDWS` parameter). This should be set to “true” thus removing the need for independent forward and backward runs.

### Appendix B.2 Parsing and Data Analysis

In this tutorial, we have recommended using a Jupyter notebook for analysis.

After initial reading and parsing, each window’s equilibration period will be automatically detected and the samples decorrelated as implemented by PyMBAR [20]. The free energies will be calculated and you will see the estimated Δ*G* with Alchemlyb-estimated statistical error. We have used conservative FEP settings in this tutorial which (though not the most efficient) should result in good convergence for this system. As noted in the previous section, more complicated systems (with more internal degrees of freedom) may require much longer sampling and narrower λ windows. In such systems, errors as high as 1 kcal/mol are not uncommon. Error estimates larger than 1 kcal/mol often indicate poor convergence, which can also be diagnosed by analyzing hysteresis. See Appendix F for more information on how to identify and resolve the underlying causes.

### Appendix B.3 Interpreting the Figures

In this section, we describe the contents and meaning of each of the figures generated by the provided Jupyter notebook. See Appendix F for strategies to address discrepancies between your own results and those described here. An example of a well-converged calculation is shown in Figure 6.

**Figure 6.**
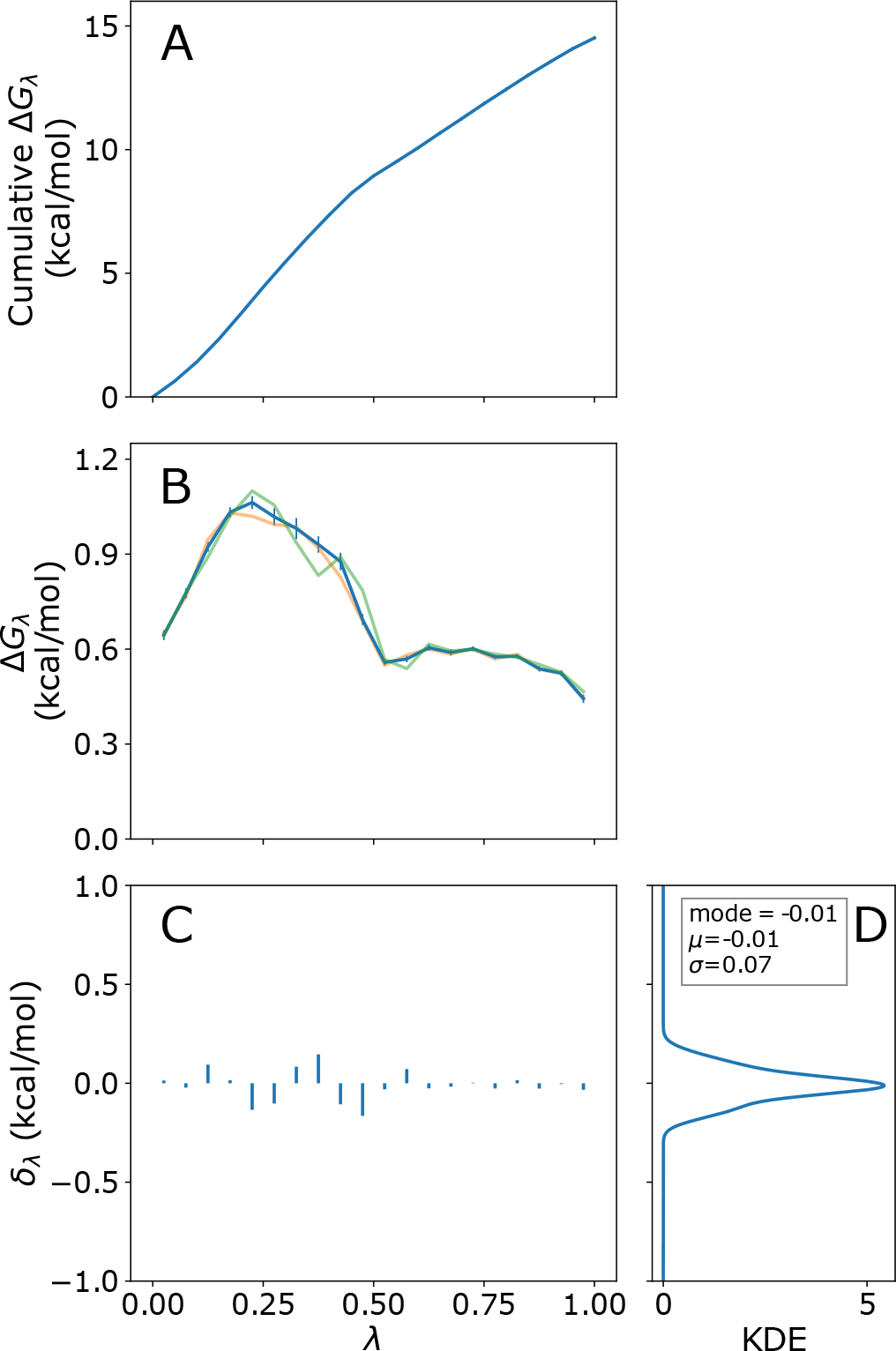
Example results from protein-phenol decoupling calculation. A) Cumulative change in free energy with accumulated error. B) Per-window difference in free energy (Δ*G*_*λ*_) calculated by the BAR estimator (blue), and exponential estimators for forward (orange) and backward (green) samples. C) Free energy hysteresis (*δ*_*λ,i*_). D) Probability density function of *δ*_*λ,i*_ estimated by kernel density estimation (KDE). Error bars indicate uncertainty reported by alchemlyb [20].

Cumulative and per-window Δ*G* curves (Figure 6 A and B) should be fairly smooth, and a magnitude of Δ*G* greater than a few kcal/mol per window is often associated with poor convergence.

For values of λ that equal `alchElecLambdaStart` or `alchVdwLambdaEnd`, a jump or cusp may be observed (such as the cusp in panel B at λ = 0.5), but discontinuities at other values of λ often indicate either insufficient samples or λ windows that are too wide.

We define hysteresis, δ_*λ,i*_, as

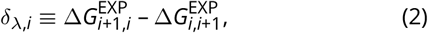

where 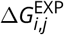 is the exponential averaging estimate of Δ*G* between λ values *i* and *j* sampled from state *i*. This is plotted in Figure 6C and D for each λ value. No value of δ_*λ,i*_ should be more than about 1 kcal/mol with a mean (μ) and mode close to zero. Similarly, the standard deviation (σ) should be less than ≈ 0.5. Failure to meet any of these criteria indicates that one or more of the λ windows has not reached equilibrium or converged.

Finally, the convergence plot should display two curves that meet quickly (before 0.5), and both curves should level out well before 1 like the example shown in Figure 7. If they are still changing at 1 or have not gotten to within 0.5 kcal/mol by 0.5, the system is unlikely to be converged at one or more λ values and the final Δ*G* estimate is likely to be inaccurate.

**Figure 7.**
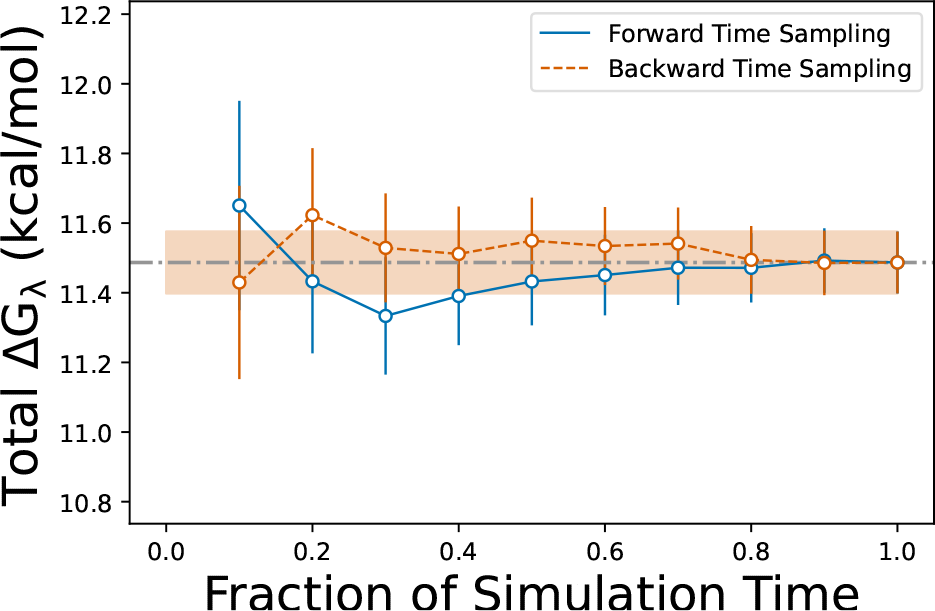
Example convergence data. We believe this calculation is well-converged due to the overlap near the halfway point and the leveling out of both curves well before the end. Error bars indicate uncertainty reported by alchemlyb.

**Figure 8.**
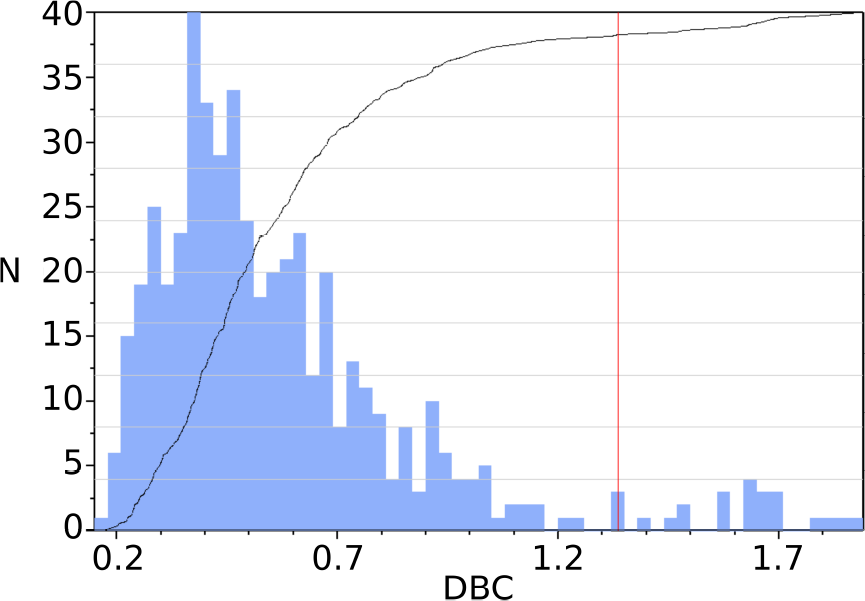
Example asymmetric DBC distribution. Screenshot from the Colvars Dashboard showing an example asymmetric DBC distribution from an unbiased simulation. Black line: cumulative distribution. Grey horizontal lines: deciles. If the phenol had flipped about the symmetric axis, there would be a second peak about 1.8 Å as seen in Fig. 10.

## Appendix C Restraints

In the simulation in which the ligand is decoupled from the site, restraints that keep the ligand from diffusing away must be applied. This serves two purposes: first, it prevents spontaneous unbinding of the ligand over the course of the simulation; second, it accelerates convergence of the computation by limiting the space to be sampled [5]. Thus restraints on the ligand are essential both for estimating a well-defined free energy of binding, and for minimizing the statistical noise on that estimate. This is often achieved by layering several rotational and translational restraints on the ligand [1, 3, 4, 6, 8, 9]. The main draw-back of such approaches is that each restraint must be designed and parameterized which adds to 1) the time and effort required to setup the simulations and 2) the complexity of the simulations themselves. As a result, troubleshooting and interpretation are more difficult and time-consuming. SAFEP in contrast uses just one restraint, the distance-to-bound-configuration (DBC), which is both robust and requires minimal parameterization [11].

### Appendix C.1 Flat-well Restraints

An ideal restraint for decoupling simulations would precisely encapsulate the bound ensemble without modifying it. That is, it would be of the form

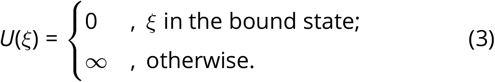

This infinitely sharp potential, however, would create numerical instability in a molecular dynamics simulation. We therefore impose continuous flat-well restraints which result in finite restorative forces when the system approaches the boundary of the bound state without modifying the bound ensemble itself. Such restraints approximate square wells with the form

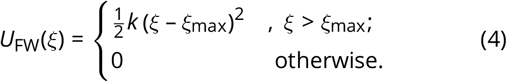

### Appendix C.2 The Distance-to-Bound Configuration (DBC) Coordinate

#### Appendix C.2.1 Definition of DBC

The DBC is the root-mean-square deviation (RMSD) of a subset of ligand atom coordinates from a typical bound pose *in the frame of reference of the binding site*. This single, scalar coordinate is designed to optimize the convergence of alchemical decoupling from the site without biasing the initial, coupled state. It captures any relative motion of the ligand with respect to the binding site as well as conformational changes of the selected ligand atoms. In general, all heavy atoms of the ligand can be included in the DBC definition, but larger, more flexible ligands may be better restrained using a smaller subset of atoms.

To impose a DBC restraint, we apply a flat-well potential defined by Equation 4 to a DBC coordinate. See Ref. 11 for details.

#### Appendix C.2.2 Symmetric DBC

One peculiarity of the model system used here is that the ligand, phenol, is symmetric. Although this isn’t strictly problematic, it does require a little extra accounting and care. This is especially true for ligands with higher symmetry numbers which would require much more complicated analytical corrections. The Colvars keyword `atomPermutation` can be used to define such symmetries:

1. The easiest way to identify equivalent atoms is to label them.
2. Use the *Graphics*→*Representations* interface to hide all atoms except the ligand.
3. Use the labeling tool to label the four symmetric carbons as shown:

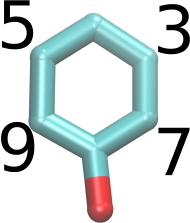
4. Open *Graphics*→*Labels*….
5. Select all four labels from the list by clicking and dragging.
6. Open the tab titled *Properties* and change the format string to %1i .
7. You may wish to adjust other settings in this menu to make the labels more visible.
8. Your view should now look something like the image above with the serial number of each atom indicated.
9. In the Colvars config editor window, place your cursor on the line before `atoms {`and add `atomPermutation {11 7 3 1 5 9 12}` .
10. Your console should now look like this:

~~~
colvar {
  name DBC_sym
  rmsd {
     atomPermutation {11 7 3 1 5 9 12}
     atoms {
        atomNumbers {11 9 5 1 3 7 12}
…
~~~

These changes make atom 7 equivalent to atom 9 and atom 3 equivalent to atom 5 for purposes of RMSD calculation.

### Appendix C.3 Isotropic center-of-mass restraint

The center-of-mass (COM) restraint is used as the container into which the ligand is released during RFEP (Step C). It is created by using the flat-well restraint in Equation 4, where ξ is the displacement of the ligand’s center of mass. The free energy cost of imposing the COM restraint can be calculated analytically because we treat the ligand as a point particle in a well-defined volume (i.e. as an ideal gas). The free energy difference between the simulated volume and an arbitrary concentration *L* is

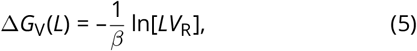

where 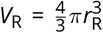 is the volume of a sphere of radius *r*_*R*_ (the upper boundary of the COM restraint) [11]. Recall that the width of the restraint is slightly larger (1 Å) than the width of the DBC restraint to avoid any edge cases in which the DBC may be larger than the COM displacement.

## Appendix D Restraint free energy calculation

### Appendix D.1 Restraint perturbation simulation

Although the DBC restraint by design does not affect the coupled state, it does modify the decoupled state and this contribution must be accounted for in the overall free energy estimation. To that end we use Colvars to run a simulation in which the DBC restraint is removed progressively and the free energy is computed for that process. To make this computation more efficient the ligand is not released into the whole simulation box, but it is kept confined in a spherical volume *V*_R_. Be advised: MD simulation algorithms can prevent center-of-mass diffusion for the whole system (in NAMD, `zeroMomentum`). In RFEP the ligand must be allowed to diffuse, so this option must be disabled.

### Appendix D.2 Thermodynamic Integration and Analysis

As in FEP, restraint free energy perturbation (RFEP) scales certain energy terms and the associated forces using a perturbation parameter, λ ∈ {0, 1}. FEP estimates finite free energy differences between λ values while TI estimates the free energy derivative. As derived and discussed in most statistical mechanics and molecular dynamics textbooks (e.g. Ref. 25, 26), the Helmholtz free energy *F* approximates the Gibbs free energy *G* when *P*Δ*V* can be neglected. Furthermore, the *P*Δ*V* terms tend to cancel out when subtracting the two alchemical free energy differences in our thermodynamic cycle, making the following approximation quite accurate:

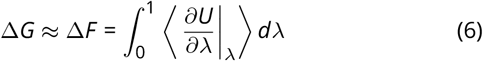

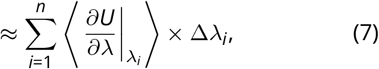

where Δλ_*i*_ is the difference between adjacent λ values.

The core of this expression, 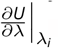, can be calculated analytically for a given value of ξ by applying the definition *k*_*λ*_ ≡ λ^*α*^ *k*_1_ – *k*_0_ to equation 4 and taking the partial derivative with respect to λ, giving

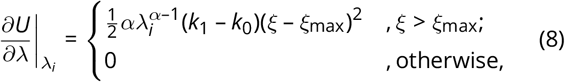

where *k*_0_ is the force constant (forceConstant) when λ = 0, *k*_1_ is the force constant when λ = 1 (targetForceConstant), and α (targetForceExponent) is a tuning parameter that improves convergence of TI by making the energy a smoother function of λ near λ = 0.

In Step C, ξ is replaced by *d*, the DBC, and ξ_max_ is replaced by the upper wall of the DBC restraint, *d*_max_. This is applied to the colvars trajectory data in the Jupyter notebook section associated with Step C (Figure 9).

**Figure 9.**
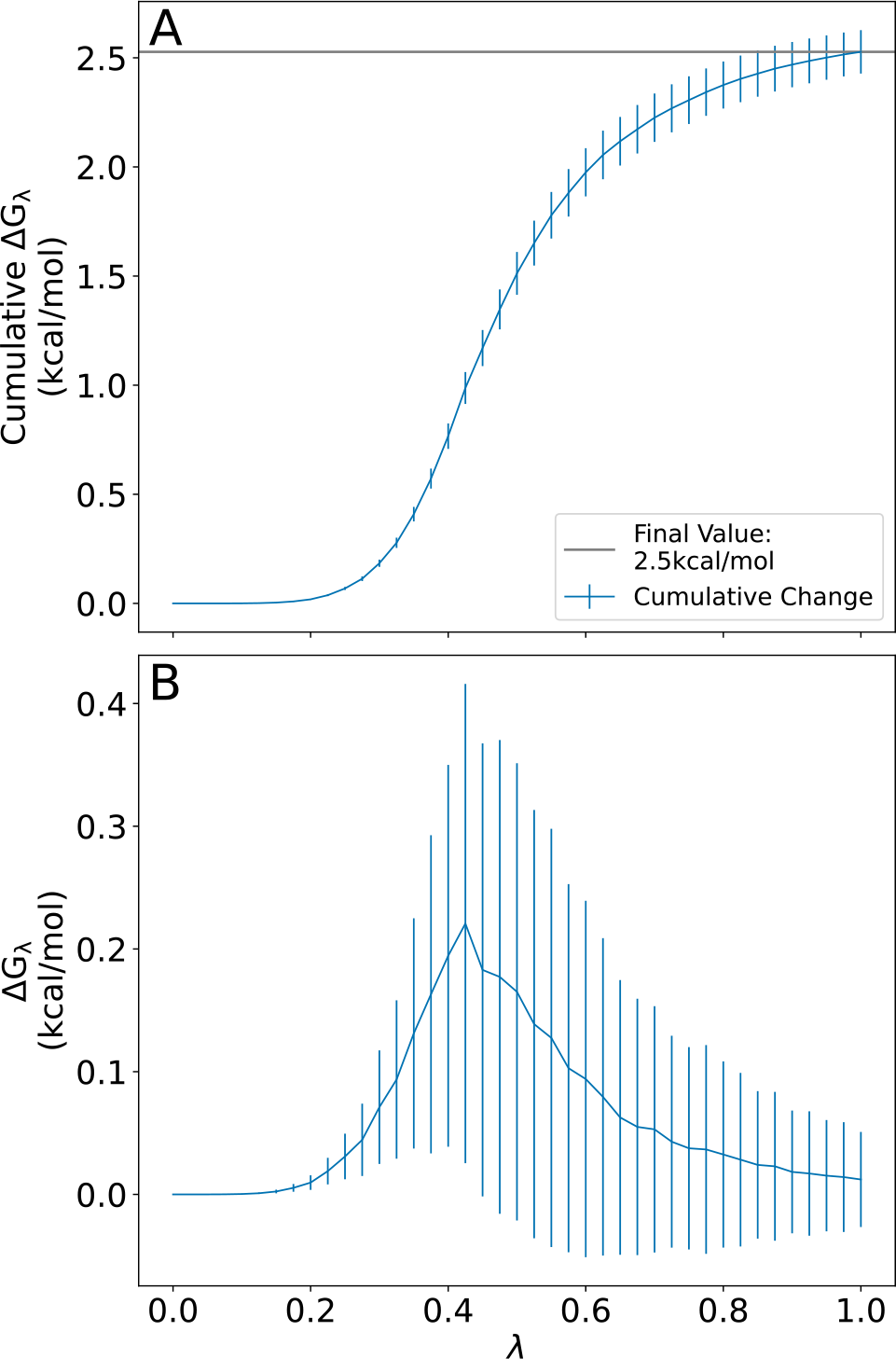
Free energy change over the course of a TI calculation. A) Restraint free energy (Δ*G*_*λ*_). B) Derivative of the free energy with respect to the coupling parameter, *λ*. Error bars indicate standard deviation.

Finally, the error is estimated by

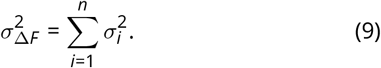

The error on each local average 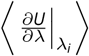 is estimated as its standard deviation, thus avoiding any assumption on the number of independent samples, leading to

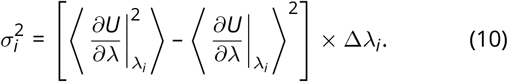

A finer estimate of the error can be obtained by running replicas of the TI calculation and computing their dispersion. Technical details of our implementation can be seen in *common/TI*.*tcl* which is just syntactic sugar for the Colvars implementation (See the [Colvars user guide](http://colvars.github.io/colvars-refman-vmd/colvars-refman-vmd.html#sec:colvarbias_ti)).

## Appendix E Concentration Dependence and Non-Ideality

While in this tutorial we have only used a single, infinitely dilute concentration to calculate Δ*G*_bind_, SAFEP can also be used to predict concentration dependence in non-ideal and non-dilute solutions. Here we consider the underlying theory for interpreting such a calculation and provide general suggestions for implementation.

We consider a “unitary” (single protein, single site [11]) system with *m* ligands in a volume *V*. The ligand concentration in this system is *L* = *m*/*V* . A DBC coordinate *d*_*i*_ can be defined for each of the *m* ligands, indexed by *i*.

The threshold on the DBC coordinate meaningfully divides the ensemble into two possible macrostates: occupied (one ligand occupies the site and *m* – 1 ligands are in solution) and unoccupied (no ligands occupy the site and *m* ligands are in solution). We formalize this here through the DBC “test function,” which for an individual ligand *i* is a Heaviside step function of the form

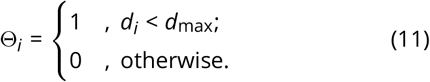

The instantaneous site occupancy Θ_occ_ is determined by whether any of the *m* ligands occupy the site, given by the sum of all the individual test functions,

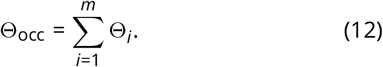

Since here we consider the case in which the site can bind at most one ligand, Θ_occ_ is either 0 or 1.

The partition functions for the occupied and unoccupied states are thus *Z*_occ_ and *Z*_*unocc*_ respectively, where

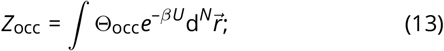

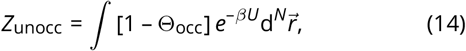

and the potential energy *U* is a function of the positions 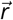 of all *N* particles in the system. Θ_occ_ is a function of the DBC coordinates *d* (and thus the positions of ligand and site atoms only).

The occupancy probability *P*_occ_(*L*) is thus

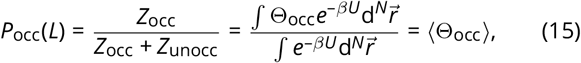

which yields the average occupancy ⟨Θ_occ_⟩.

*G*_occ_ and *G*_unocc_ are the free energies of the occupied and unoccupied macrostates respectively, where

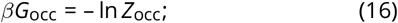

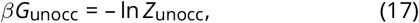

so

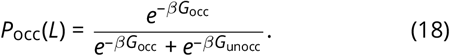

We turn now to connecting these quantities to a SAFEP calculation. In step D of the protocol, we decoupled one ligand from a bulk that contained *m* = 0 fully coupled ligands, for an infinitely dilute concentration of *L* = 0/*V* . We then extrapolated to the standard concentration using an ideal gas correction that assumes ideality.

For a ligand at finite concentration in a non-ideal bulk, it is neither useful or necessary to standardize the free energy. Instead, we would carry out Step D at the finite ligand concentrations of interest (*L* = *m*/*V* > 0), and adjust Step E to calculate the unstandardized free energy Δ*G*_bind_(*L*) as follows:

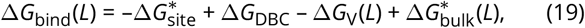

where the volume per molecule in the bulk is *V* /*m* and thus

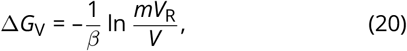

since Δ*G*_bind_(*L*) = *G*_occ_(*L*) – *G*_unocc_(*L*).

Equation 18 can be rewritten in terms of Δ*G*_bind_(*L*),

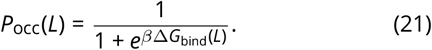

Even for a non-ideal bulk, we may assume the excess chemical potential is unchanging for small changes in concentration. Thus we would perform ligand decoupling (step D) at finite concentration *L*, and use the ideal gas correction to predict occupancy for nearby concentrations *L*^′^, as long as |*L*–*L*^′^| is small,

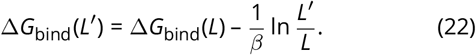

Substitution of Equation 22 in Equation 21 yields the occupancy probability for concentration *L*^′^,

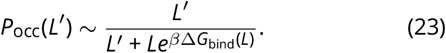

Incidentally, for dilute *L*, 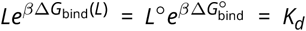, and Equation 23 reduces to a form familiar to biochemists 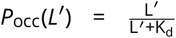. In general, Equation 22 only holds if the change in excess chemical potential is negligible between *L*^′^ and the simulation concentration *L*. This assumption can be tested by running bulk decoupling (Step D) at both *L*^′^ and *L* and checking that the resulting change in Δ*G*_bulk_ is much smaller than the overall error. If we wish to calculate *P*_occ_ over a wider concentration range in which this assumption does not hold, we would need to explicitly calculate 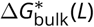 for multiple simulation concentrations *L* and extrapolate to the intermediate concentrations, as in Ref. 11.

## Appendix F Troubleshooting

We have written this tutorial to be as robust as possible but also generalizable to other systems. In the process of applying these steps to your own system of interest, however, additional challenges may arise. When calculations fail to converge or appear to converge to unreasonable values, it can be difficult to discern what has gone wrong without simply starting over. We provide here some of the most common issues and their respective fingerprints as cautionary tales and troubleshooting tools. If you encounter a problem with running the tutorial as written and do not see your issue listed below, please contact us.

### Appendix F.1 Problems with Running; NAMD crashes

Alchemical FEP will make any instabilities in a system more apparent as well as introduce a few more possible sources of instability. The most common problems are RATTLE errors and box size instability.

#### Appendix F.1.1 RATTLE Errors

~~~
ERROR: Constraint failure in RATTLE
algorithm for atom 593!
~~~

**Causes:**

If the usual culprits (poor equilibration, long time steps, and over-aggressive RESPA settings) have been ruled out, the most likely causes of RATTLE errors during FEP are 1) too-wide λ windows and 2) a too-low soft-core potential exponent.

**Solutions:**

Lambda windows can easily be narrowed by reducing dLambda. Values between 0.05 and 0.005 give a good balance between efficiency and accuracy. Note, decreasing dLambda will result in more windows which will require more CPU time overall.

#### Appendix F.1.2 Box Size Instability

~~~
FATAL ERROR: Periodic cell has become
too small for original patch grid!
Possible solutions are to restart
from a recent checkpoint, increase
margin, or disable useFlexibleCell
for liquid simulation.
~~~

**Causes:**

While classical MD is “tolerant” to small periodic boxes and aggressive barostats, combining these with FEP is particularly unstable.

**Solutions:**

First, the periodic box should be at least twice the solute size or twice the cutoff distance (whichever is longer) in order to avoid violating the minimum image convention which can cause instabilities during FEP decoupling, especially with charged solutes. Second, at least with the Langevin barostat, slowing the piston dynamics can improve system stability at the cost of slowing box-size relaxation. We use LangevinPistonPeriod between 75 and 200, and LangevinPistonDecay between 50 and 100. LangevinPistonDecay should always be about half LangevinPistonPeriod. See [The NAMD UG](https://www.ks.uiuc.edu/Research/namd/2.14/ug/node39.html) for more details [13].

### Appendix F.2 Problems with Results; Poor Convergence

Convergence within each step is a prerequisite to a reliable final estimate of 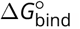. Large errors and internal inconsistencies often indicate poor equilibration or under-sampling of one or more ensembles. Each leg of a SAFEP calculation has unique challenges and edge-cases which we address below. In general, convergence may be improved by increasing the simulation time for each λ value.

#### Appendix F.2.1 Local and Misleading Convergence

A FEP calculation may converge locally and give a biased outcome, in the case of very slow fluctuations reflecting the presence of metastable states. The best way to detect this is to run multiple replicas that are as weakly correlated as possible (i.e. true replicas are better than pseudo-replicas with identical initial conditions, and pseudo-replicas are better than no replicas).

The analysis of the protein-ligand bound state ensemble is required because it directly affects the definition of the DBC. Simulation of the apo protein (without ligand in the binding pocket), however, can also provide useful information about the decoupled endpoint. In the case of lysozyme, for example, the binding pocket is frequently occupied by one or two water molecules. If the lysozyme binding pocket does not recover hydration once the ligand is fully decoupled during FEP, the calculation overestimates the strength of binding by up to 0.5 kcal/mol. This is a small error compared to the overall precision of the technique, but users should be aware that assessing endpoint hydration is particularly important for larger or more hydrated binding pockets.

#### Appendix F.2.2 Step A: Define the DBC

**Symptom:** Multimodal DBC distribution (e.g. Figure 10).

**Figure 10.**
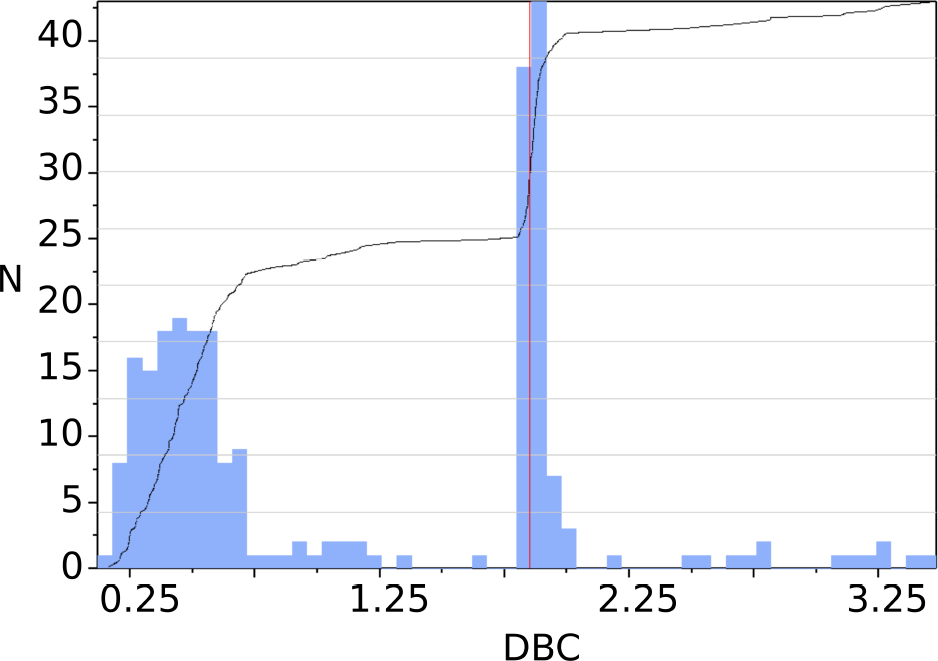
Screenshot from the Colvars Dashboard showing a bimodal distribution that resulted from using an asymmetric DBC for phenol. The second peak corresponds to a 180 degree rotation about the C-O axis.

**Causes:**

There are three main causes of multimodal DBC distributions: 1) ligand unbinding, 2) multiple binding modes, and 3) multiple nearby binding sites.

**Solutions:**

Use the Colvars Dashboard histogram tool to probe the conformations associated with each mode and decide which modes correspond to bound and unbound states.

|  |  |
| --- | --- |
| Bound and Unbound modes | If all but one mode may be described as unbound, place the DBC restraint upper wall between the bound mode and the unbound mode with the lowest DBC value. Proceed with FEP. |
| Multiple, indistinguishable bound modes | Such modes are a result of symmetric ligands and are best addressed using a symmetric DBC. See Appendix C.2.2 for more details. |
| Multiple, distinguishable bound modes | If one or more bound mode(s) is meaningfully distinct from some other mode(s), select a representative frame for each class. These frames become reference poses for each binding “site” from which you must calculate $\Delta G_{\text{site}}$, $\Delta G_{\text{DBC}}$, and $\Delta G_v$ separately. Free energy contributions from each mode can be combined by exponential summing as shown in equation 12 of [27]. |

#### Appendix F.2.3 Step B or D: FEP Calculations

**Symptom:** DBC restraint energy stays close to 0.

**Causes:**

The DBC restraint may be too wide (upperWalls) or too soft (forceConstant). The λ windows may be too short to properly sample the decoupled ensemble.

**Solutions:**

First, watch the trajectory for any abnormal behavior; wide DBCs will often be obvious from the last several nanoseconds of a FEP run because the ligand will explore a much larger conformation space than expected. See the DBC debugging checklist below. If the system passes all checks, try running λ = 1 for longer to make sure the DBC restraint is functioning.

**Symptom:** DBC energy is consistently greater than 0.

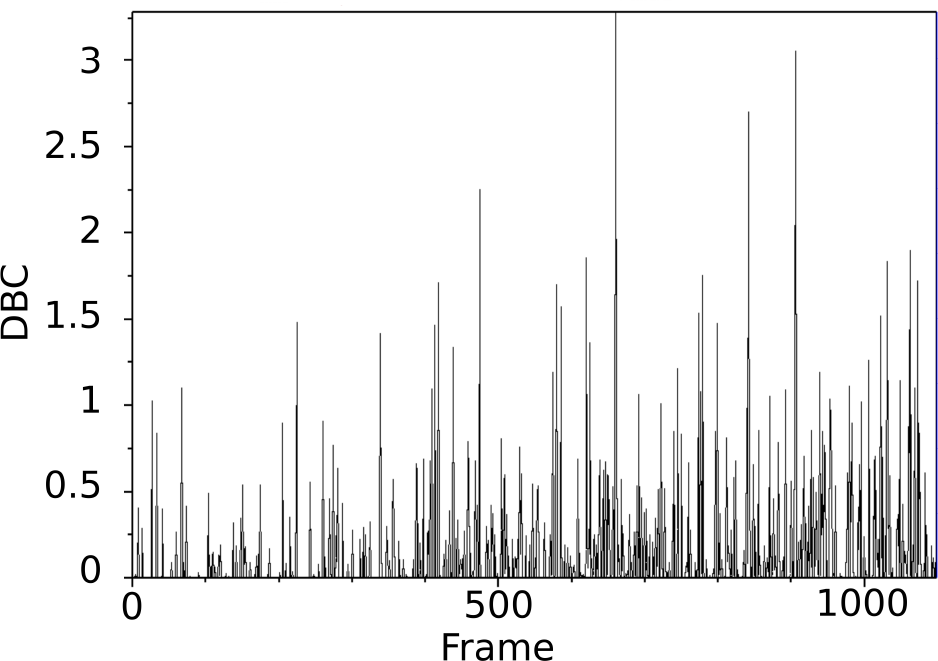

**Causes:**

This issue is most often due to too-narrow DBC restraints (upperWalls) or other mistakes in the colvar definition.

**Solutions:**

This problem is harder to diagnose from the trajectory alone unless there are obviously over-restrained degrees of freedom in the ligand. Consult the DBC debugging checklist below.

**Symptom:** Ligand unbinds during FEP.

**Cause:**

The most likely cause of unbinding during FEP is a DBC restraint that is too wide (upperWalls), too soft (forceConstant), inactive, or improperly defined.

**Solutions:**

Consult the checklist below:

**DBC Debugging Checklist:**

□ Only the DBC restraint should be active during FEP (Step B)
□ DBC restraint upper walls have the intended value. (3.b)
□ DBC restraint force constant is appropriate (100 or 200). (3.b)
□ NO lower walls (3.b)
□ If the Colvars configuration file contains a “width” keyword, it should be 1. See [28] and the [Colvars user guide](http://colvars.github.io/colvars-refman-vmd/colvars-refman-vmd.html#sec:colvar_grid_params) for more details. (3.b)

**Symptom:** Hysteresis larger than 1 kcal/mol for any λ window.

**Causes:**

Large hysteresis values are most often caused by: 1) insufficient equilibration, 2) short windows (less than a few hundred ps), or 3) wide windows (large dLambda).

**Solutions:**

If the system is well-relaxed and equilibrated by the usual metrics (box size, pressure, temperature, etc.), then it is most likely that either the λ windows are too short or too wide. Try increasing the sampling time or increasing the total number of windows. We have had good results with 120 windows of 3 ns each, but longer may be necessary for particularly unwieldy systems.

**Symptom:** Very large hysteresis near λ = 0 or λ = 1.

**Causes:**

Large hysteresis near the end-points of the FEP calculation are most commonly caused by so-called “end-point catastrophes.” See [The NAMD UG](https://www.ks.uiuc.edu/Research/namd/2.14/ug/node63.html) for more details [13].

**Solutions:**

Ensure alchVdwShiftCoeff is between 5 and 8. If this is already the case, and no other part of the calculation is problematic, try doubling the number of windows between the window with large hysteresis and the nearest end-point.

**Symptom:** Hysteresis oscillates or is otherwise correlated with λ.

**Causes:**

1. Some versions of NAMD have a bug that allows FEP data to be written on a step without energy calculations. This results in the use of stale energies (from a previous step) and inaccurate estimates for differences in energy.

2. The λ windows may be too short to reach equilibrium

**Solutions:**

As noted above, δ_*λ,i*_ should be independent of λ with a mean of 0.

1. Manually ensure that alchOutFreq is a multiple of both fullElectFrequency and nonbondedFreq. See [Bug advisory and Workaround](https://www.ks.uiuc.edu/Research/namd/mailing_list/namd-l.2020-2021/1487.html) for more details.

2. Extend the length of each window or decrease dLambda.

#### Appendix F.2.4 Step C: TI Calculation

The DBC restraint should do the most work early in the TI calculation then, as the force constant is scaled out, the COM restraint should take over and keep the center of mass in a well-defined volume. TI convergence issues are most easily diagnosed by watching the MD trajectory and examining the colvar trajectories.

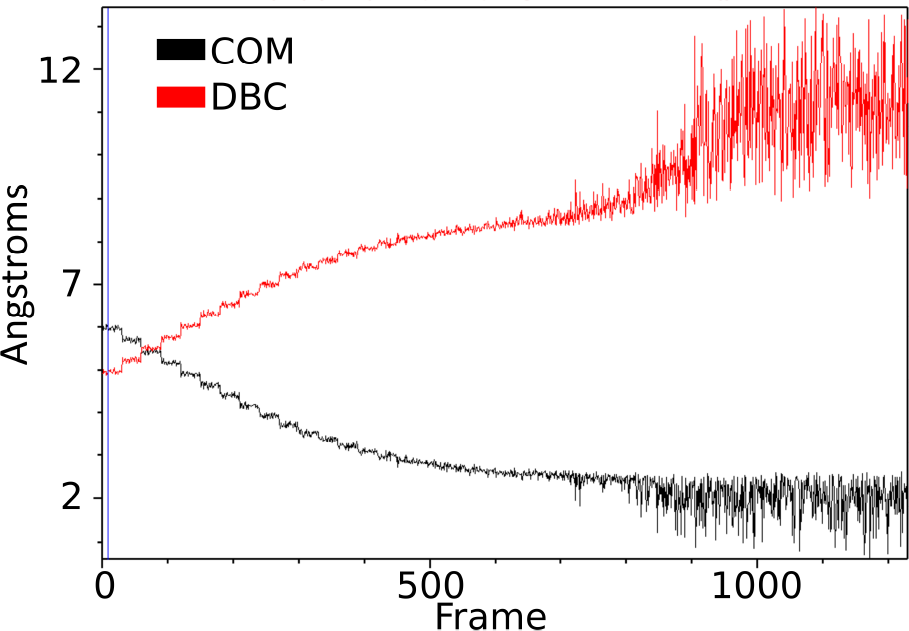

**Symptom:** The collective variable trajectory is abnormal.

**Causes:**

Strange behavior is often caused by a mismatch between the coordinates used to determine the center of the COM restraint and the reference coordinates used for the DBC restraint as is the case in the figure above.

**Solutions:**

Update the reference coordinates to match those used for determining the center of the COM restraint. That is, revisit Step C1.b paying special attention to the coordinates used. Make sure you save your refPositionsFile paths are correct.

**Symptom:** COM restraint energies during TI are very high or very low and don’t change much.

**Causes:**

The harmonic walls forceConstant is likely too stiff (high) or too soft (low).

**Solutions:**

Adjust the force constants to be between 100 (minimum) and 200.

**Symptom:** The ligand doesn’t move during TI (or moves very little).

**Causes:**

1. The COM restraint is too narrow (upperWalls is too small).
2. The run is too short to visit all λ values.
3. The DBC restraint isn’t being released.

**Solutions:**

1. Double check the choice of wall position and ensure that it is correct in all Colvars configuration files.
2. Make sure your run is long enough; if you increase the targetNumSteps parameter, you must also increase the run time proportionally.
3. If you are using our *TI*.*tcl* script, double-check all settings and parameters. If you have written your own TI Colvars config file, check all settings and compare them to the settings reported in the NAMD log file.

**Symptom:**

The TI calculation doesn’t converge *and* the ligand moves very far away from its initial position.

**Causes:**

The COM restraint is probably too wide (or non-existent).

**Solutions:**

Double-check the existence and parameters of the COM restraint.

## 4 Author Contributions

ESM, ME and JWS ran, tested and refined the protocol. ESM wrote the notebook and analysis scripts. GB and JH designed and supervised the work. All authors wrote the document.

## 5 Other Contributions

The authors are grateful to Ms. Noureen Abdelrahman, Ms. Mariadelia Argüello-Acuña, Mr. Jahmal Ennis, Mr. Connor Pitman, and Mr. Jules Marien for providing feedback and initial testing of this tutorial. The authors acknowledge the Office of Advanced Research Computing (OARC) at Rutgers, The State University of New Jersey for providing access to the Amarel cluster and associated research computing resources.

## 6 Potentially Conflicting Interests

The authors declare no potential conflict of interest.

## 7 Funding Information

We acknowledge financial support from the National Science Foundation DGE 2152059 (to ESM, JWS, and GB), the Ministry of Science, Research, and Technology of the Islamic Republic of Iran for a Ph.D. candidate research grant to ME, and the French Agence Nationale de la Recherche (ANR) for grant LABEX DYNAMO (ANR-11-LABX-0011-01, to JH).

## Notes

https://github.com/jhenin/SAFEP_tutorial

### Competing Interest Statement

The authors have declared no competing interest.

### Summary of Updates

Typographic and punctuation corrections Layout improvements Miscellaneous clarifications

